# TAILS identifies candidate substrates and biomarkers of ADAMTS7, a therapeutic protease target in coronary artery disease

**DOI:** 10.1101/2021.11.19.469331

**Authors:** Bryan T. MacDonald, Hasmik Keshishian, Charles C. Mundorff, Alessandro Arduini, Daniel Lai, Kayla Bendinelli, Nicholas R. Popp, Bidur Bhandary, Karl R. Clauser, Harrison Specht, Nadine H. Elowe, Dylan Laprise, Yi Xing, Virendar K. Kaushik, Steven A. Carr, Patrick T. Ellinor

**Affiliations:** Cardiovascular Disease Initiative, Broad Institute of MIT and Harvard, Cambridge, MA; Proteomics Platform, Broad Institute of MIT and Harvard, Cambridge, MA; Center for the Development of Therapeutics, Broad Institute of MIT and Harvard, Cambridge, MA

## Abstract

Loss-of-function mutations in the secreted enzyme ADAMTS7 (a disintegrin and metalloproteinase with thrombospondin motifs 7) are associated with protection for coronary artery disease (CAD). ADAMTS7 catalytic inhibition has been proposed as a therapeutic strategy for treating CAD; however, the lack of an endogenous substrate has hindered the development of activity-based biomarkers. To identify ADAMTS7 extracellular substrates and their cleavage sites relevant to vascular disease, we used TAILS (terminal amine isotopic labeling of substrates), a method for identifying protease-generated neo–N termini. We compared the secreted proteome of vascular smooth muscle and endothelial cells expressing either full-length mouse ADAMTS7 WT, catalytic mutant ADAMTS7 E373Q or a control luciferase adenovirus. Significantly enriched N-terminal cleavage sites in ADAMTS7 WT samples were compared to the negative control conditions and filtered for stringency, resulting in catalogs of high confidence candidate ADAMTS7 cleavage sites from our three independent TAILS experiments. Within the overlap of these discovery sets, we identified 24 unique cleavage sites from 16 protein substrates, including cleavage sites in EFEMP1 (EGF-containing fibulin-like extracellular matrix protein 1/Fibulin-3). The ADAMTS7 TAILS preference for EMEMP1 cleavage at the amino acids 123.124 over the adjacent 124.125 site was validated using both endogenous EFEMP1 and purified EFEMP1 in a binary *in vitro* cleavage assay. Collectively our TAILS discovery experiments have uncovered hundreds of potential substrates and cleavage sites to explore disease related biological substrates and facilitate activity-based ADAMTS7 biomarker development.

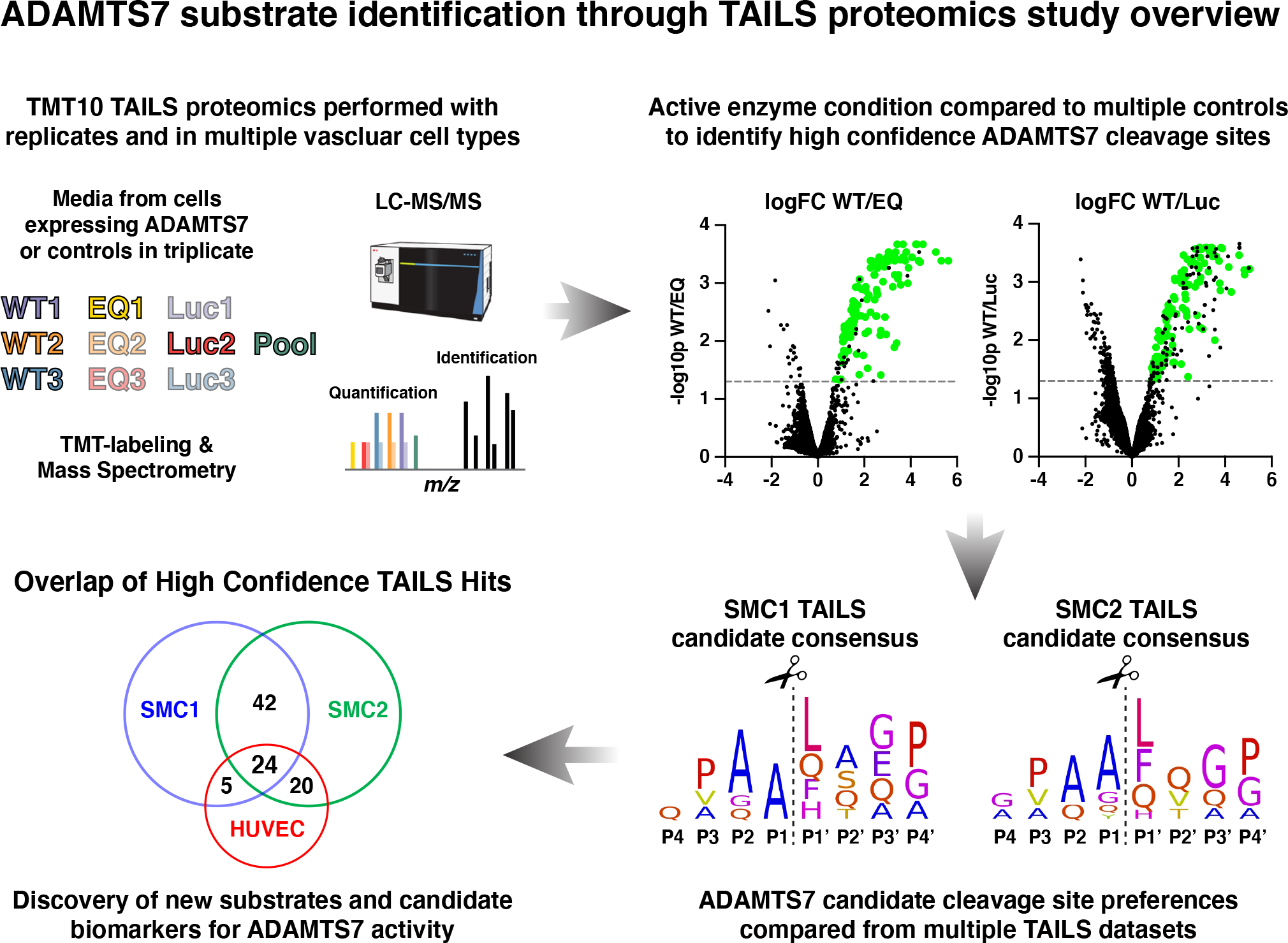
Graphical Abstract.

## Introduction

ADAMTS7 is a large, secreted protease with poorly defined substrates. Variants at the *ADAMTS7* locus were identified in population based genome wide association studies (GWAS) for CAD, with a risk haplotype including a coding variant in the prodomain, rs3825807 p.Ser214Pro (1-3). The Ser214 risk variant was shown to increase prodomain processing and maturation, correlating with an increase in COMP (Cartilage oligomeric matrix protein) degradation and vascular smooth muscle cell migration (4). Additionally, the Ser214 risk allele was associated with an unstable atherosclerotic plaque phenotype and an increase in secondary cardiac events (5-8). Mouse knockout studies have shown that *Adamts7* loss-of-function significantly reduces vascular smooth muscle cell mediated neointimal formation in arterial wire injury models (9, 10). These findings were supported by *Adamts7* knockdown experiments in a rat carotid artery balloon injury model of restenosis (11, 12). Conversely, ADAMTS7 overexpression in the rat balloon injury model increased the rate of neointima formation, presumably through an increase in extracellular matrix degradation to allow for vascular smooth muscle cell migration at the site of vascular injury. ADAMTS7, and its paralog ADAMTS12, are defined as COMP proteases that utilize their carboxyl terminal regions for substrate recognition (13, 14). Based on evidence from gel-based cleavage assays, the carboxyl terminal region of ADAMTS7 was shown to be necessary for substrate cleavage, demonstrating a requirement of the full-length ADAMTS7 protease (15). However, the ADAMTS7 substrate cleavage sites for COMP and several other reported substrates from gel-based assays have not been identified. Furthermore, the lack of an endogenous substrate cleavage site for ADAMTS7 has hindered the development of activity-based biomarkers for clinical development.

ADAMTS7 belongs to a family of 19 secreted zinc metalloproteinases with a shared organization of a signal peptide, prodomain, metalloproteinase, disintegrin, thrombospondin, cysteine-rich and spacer domains (16). Additionally, ADAMTS7 has a total of eight thrombospondin type I repeats and a highly glycosylated mucin domain with a chondroitin sulfate glycosaminoglycan (CS-GAG) attachment that set ADAMTS7 and ADAMTS12 apart from their family members (17, 18). Consequently, the CS-GAG modified ADAMTS7 is both an extracellular protease and a proteoglycan. Other ADAMTS family members have known substrates and are associated with human disease (19). *ADAMTS13* loss-of-function mutations or auto-immune function blocking antibodies to ADAMTS13 result in the clotting disorder thrombotic thrombocytopenic purpura (TPP). During hemostasis, von Willebrand factor (vWF) is precisely cleaved by ADAMTS13, using an elaborate substrate recognition system involving the disintegrin, cysteine-rich and spacer domains (20). Loss-of-function mutations in the procollagen I N-proteinase *ADAMTS2* result in the connective tissue disorder dermatosparaxis Ehlers-Danlos syndrome (dEDS). ADAMTS4 and ADAMTS5 (also known as aggrecanase-1 and aggrecanase-2) degrade the extracellular matrix proteins aggrecan and versican at defined cleavage sites. Antibodies raised to the aggrecan neo-epitopes are in use for clinical development of osteoarthritis therapeutics targeting ADAMTS5 enzymatic function (21, 22).

TAILS proteomics is an un-biased method for identifying cleavage sites from full-length substrates in their natural context (23). In this protocol, the primary amine groups from samples with and without protease activity are labeled and blocked by isobaric tags. N-terminally labeled peptides with higher ratios in the enzyme treated samples represent neo N-termini or candidate cleavage sites for the enzyme. The TAILS method has been applied for the secreted metalloproteinases MMP2 and MMP9 (24) and for the procollagen I N-proteinases ADAMTS2, ADAMTS3 and ADAMTS14 (25, 26) to identify additional substrates and substrate cleavage sites. Recently TAILS was applied to ADAMTS7 using one active and one inactive condition from a truncated ADAMTS7 to identify substrates from a human fibroblast cell line, including a candidate site in LTBP4 (Latent-transforming growth factor beta-binding protein 4) (27).

Here we present the results from TAILS experiments performed in human coronary artery smooth muscle cells and vascular endothelial cells using full-length WT and catalytic mutant ADAMTS7. In three independent experiments, we identified 24 unique cleavage sites from 16 protein substrates, including cleavage sites in EFEMP1 (EGF-containing fibulin-like extracellular matrix protein 1/Fibulin-3).

### Experimental Procedures

#### ADAMTS7 Expression, Cell Culture and Preparation of the Secreted Proteins

Full-length mouse *Adamts7* 1-1857 (Uniprot Q68SA9, wild type (WT) or E373Q (EQ) catalytic mutant) with a carboxyl terminal 3xFLAG tag was used to generate custom adenoviruses as previously described (28). Adeno-CMV-Luciferase (Luc) from Vector Biolabs was used as a negative control. Human Coronary Artery Smooth Muscle Cells (HCA-SMC) were obtained from Lonza and expanded in Lonza SmGM-2 media in 15cm tissue culture plates. Twenty-four hours before media collection, near confluent plates were transduced with 50 MOI adenovirus and switched to Lonza Basal SM media with volumes of 20ml per 15cm plate. Expression of the Flag tagged ADAMTS7 was confirmed from the media using western blot and detected using the anti-Flag M2-HRP antibody (Sigma, A8592). Collected media was processed by adding protease inhibitors (1mM EDTA and 1mM PMSF) and clarified by 500g x 5min spin at 4C to pellet cell debris. Cell media supernatants were then passed through a 0.22um filter and kept chilled. Processed media was then concentrated 50-100x using 3kDa Centricon Plus-70 filter unit, spun at 3,500g x 60min at 4C. Approximately 10% of the processed, concentrated media was set aside for total proteome analysis of the secretome. The remaining 90% of the processed, concentrated media was buffer exchanged into 50mM HEPES pH 8.0, 150mM NaCl (2 x 50ml) using the 3kDa Centricon Plus-70 filter unit and concentrated to >2mg/ml to serve as input for the TAILS experiment. Samples for secretome proteome and TAILS were stored at -80°C until further use.

For the first experiment, referred to as SMC1, 120ml of media from 6x 15cm dishes was pooled from each condition (Luc, WT, EQ) and processed to produce three separate inputs for the TAILS experiment. Luc, WT and EQ inputs for the SMC1 experiment were each divided into triplicate 400ug samples to generate technical replicate TAILS data. For the second experiment, referred to as SMC2, media was processed separately in triplicate for each condition (Luc1, Luc2, Luc3, WT1, WT2, WT3, EQ1, EQ2, EQ3) using 60ml of media from 3x 15cm dishes for each process replicate. Inputs from the SMC2 experiment used process replicates of 300ug each to generate TAILS data. The third experiment used input from cultured Human umbilical vein endothelial cells (HUVEC) from Lifeline Cell Technology grown in VascuLife Media and transduced in Lifeline Basal EC (+bFGF2 10ng/ml). The HUVEC experiment used the same strategy as the SMC2 experiment, with 40ml of media from 2x 15cm dishes for each process replicate and used 200ug each to generate TAILS data. Inputs for process replicate experiments were restricted to the lowest concentration from the 9 parallel samples for SMC2 or HUVEC.

#### TAILS sample preparation for LC-MS/MS analysis

Samples were prepared according to the TAILS protocol from the Overall lab (23) with some modifications. For each experiment, nine samples in 50mM HEPES pH 8.0, 150mM NaCl were first denatured with guanidinium chloride (Sigma, G4505) to reach a final concentration of 2.5M guanidinium chloride and 250mM HEPES and incubated at 65⁰C for 15min. Reduction was achieved through addition of tris(2-carboxylethyl)phosphine (TCEP) at a final concentration of 20nM TCEP and incubated at 65⁰C for 45min. Alkylation reaction was performed with addition of iodoacetamide (IAA, Sigma, A3221) to a final concentration of 10mM IAA for 15min at room temperature in the dark.

Following denaturation, reduction and alkylation steps, ten percent of each of the nine samples was removed and combined to serve as a pooled reference. Isobaric labeling at protein level with TMT10 (ThermoFisher, 90113) reagent was performed at the ratio of 10:1 TMT:protein in 50% DMSO, shaking at 850rpm. TMT channels were randomly assigned to the nine samples and TMT 131 was assigned to the pooled reference sample created by pooling equal amount from all 9 samples. A second round of TMT labeling was performed sequentially for 30min more to increase labeling efficiency. Labeling reactions were quenched with 100mM ammonium bicarbonate for 15 min at room temperature. Five percent of each reaction was removed to assess isobaric labeling efficiencies. Ten TMT labeled samples were then combined and precipitated using 8 volumes of cold acetone and 1 volume of cold methanol in Beckman BK357001 tubes and stored at -80⁰C for 3 hours. Mixed sample as well as 10 labeling efficiency samples were then centrifuged in a JA-17 rotor at 14,000g for 20min. Supernatants were discarded and the samples were washed twice with 20ml of ice-cold methanol to remove residual guandinium chloride before trypsin digestion. Pellets were air dried (with SpeedVac briefly when needed), resuspended with 50mM NaOH and adjusted to 1mg/ml protein in 50mM HEPES pH 8.0. SMC1 experiment was digested solely with sequencing grade Trypsin (Promega, V5113) at a ratio of 1:50 protease to protein at 37⁰C overnight. For SMC2 and HUVEC experiments, half of the pooled sample was digested with Trypsin and the other half was digested with sequencing grade AspN (Promga, V1621) at 37⁰C overnight, and then mixed together after quenching the reactions the next day. Five percent of each pooled digestion reaction (preTAILS) was removed to assess negative selection efficiencies. Digested samples were adjusted to pH 6-7 and were enriched for TMT blocked N-termini using a hyperbranched polyglycerol aldehyde polymer (HPG-ALD) from the Kizhakkedathu lab, University of British Columbia (Flintbox). HPG-ALD was washed with water and added at 5-fold excess to the digested protein with sodium cyanoborohydride (20mM final concentration) and incubated at 37⁰C overnight. Polymer and polymer bound peptides were retained in 3kDa Amicon column and the flow-through representing labeled peptides was collected for liquid chromatography-mass spectrometry (LC-MS/MS) analysis. Five percent of the flow-through (postTAILS) was removed to assess negative selection efficiencies. Enrichment efficiency was evaluated by comparing preTAILS and postTAILS isobaric labeling efficiencies for each experiment:

Remaining flow-through was desalted on a 30mg Oasis HLB cartridge. After sample cleanup, the flow-through was separated using basic reverse-phase chromatography on a 2.1 x 250mm Zorbax 300 extend-c18 column with a 60min gradient using 20mM Ammonium Formate/ 2% Acetonitrile (ACN) pH 10 as buffer A and 20mM Ammonium Formate / 90% ACN pH 10 as buffer B. The sample was separated into 96 fractions and concatenated down to 12 by combining every 13th fraction. The 12 fractions were dried in the SpeedVac, reconstituted in 9uL of 3%/0.1% ACN/Formic acid and 4uL of it was analyzed by LC-MS/MS

### Secretome proteome sample preparation for LC-MS/MS analysis

Sample processing for analysis of the total secretome was performed as previously described (29). Briefly, samples were reduced, alkylated and LysC/trypsin digested followed by TMT labeling of 40ug aliquots for each. Similar to the TAILS experiment, channels were assigned randomly and channel 131 was used for the pooled peptide reference consisting of equal amounts of all nine samples. After confirming over 95% labeling efficiency, the reactions were quenched, samples were mixed and desalted on 30mg Oasis HLB cartridge. Sample was then fractionated as described above and resulting 12 fractions were reconstituted in 30ul of 3%/0.1% ACN/Formic acid and 1uL of it was analyzed by LC-MS/MS.

### LC-MS/MS Analysis

The TAILS and proteome fractions were separated on a Proxeon nanoLC using 3%/0.1% ACN/FA for Buffer A and 90%/0.1% ACN/FA for Buffer B on a 27cm 75um ID picofrit column packed in-house with Reprosil C18-AQ 1.9um beads (Dr Maisch GmbH)C with a 90min gradient consisting of 6-20% Buffer B in 62min, 20-35% B for 22min, 35-60% B for 9min, 60-90% B for 1min followed by a hold at 90% B for 5min. Online LC-MS/MS was performed on a Thermo Q-Exactive Plus mass spectrometer. The MS method consisted of a full MS scan at 70,000 resolution and an AGC target of 3e6 from 300-1800 m/z followed by MS2 scans collected at 35,000 resolution with an AGC target of 5e4 with a maximum injection time of 120ms and a dynamic exclusion of 20 seconds. The isolation window used for MS2 acquisition was 0.7 m/z and 12 most abundant precursor ions were fragmented with a normalized collision energy (NCE) of 29 optimized for TMT10 data collection.

### Peptide/Protein quantification, and statistical analysis

Both TAILS and proteome datasets were processed using Spectrum Mill MS Proteomics Software. The raw MS files were extracted and searched against the Uniprot human database downloaded on December 28th 2017 with the mouse ADAMTS7 sequence appended. For TAILS datasets in the SMC1 experiment the search was done in 2 rounds while in SMC2 and HUVEC experiments with the split AspN/Trypsin digest in a total of 4 rounds. The first search using a tryptic cleavage motif (K.; R.) was done with TMT10 as a fixed modification of peptide N-termini and Lysine side chains and acetylated protein N-termini as a variable modification to identify all TMT labeled peptides and acetylated N-terminal peptides of proteins. Peptide spectra not identified in this first search were searched again adding acetylated peptide N-termini to the fixed modifications to identify any peptides that are acetylated but aren’t the N-termini of the protein. This process was then repeated using an AspN (.D; .E) cleavage motif for SMC2 and HUVEC experiments. Proteome datasets were searched once with trypsin cleavage, and TMT10 as fixed modification. Peptides were autovalidated with score optimization to achieve less than 1% false discovery rate (FDR). For the proteome datasets additional protein level autovalidation was performed with the criteria of 0% FDR at protein level and minimum protein score of 13.

Distinct peptide and protein level exports were generated in Spectrum Mill for TAILS and proteome datasets, respectively. TMT reporter ion ratio of every channel to the pooled reference channel was used for the quantification and further statistical analysis. TMT10 reporter ion intensities were corrected for isotopic impurities in the Spectrum Mill protein/peptide summary module using the afRICA correction method which implements determinant calculations according to Cramer’s Rule and correction factors obtained from the reagent manufacturer’s certificate of analysis (https://www.thermofisher.com/order/catalog/ product/90406) for lot numbers UA280170 and TE270748 for SMC and HUVEC datasets, respectively (30). In TAILS datasets TMT ratios of each sample were normalized to the median ratio of the quantified natural N-termini peptides. This was done by filtering for TMT labeled peptides at the N-termini with a start amino acid number of 1 or 2 and finding the median ratio of these peptides in each channel. A moderated two sample T-test was used to compare the different groups together. Significant regulation was assessed using Benjamini Hochberg corrected p-values, referred as adjusted p-values below. Correlation plots and heatmaps of regulated peptides/proteins was performed in Protigy (Proteomics Toolset for Integrated Data Analysis, Broad Institute).

### Candidate prioritization

From the TAILS datasets, the adjusted p-value (adj.P.val) and log Fold Change (logFC) values were used to identify substrate cleavage sites enriched for ADAMTS7 proteolytic activity. Results plotting logFC and -log10 (adj.P.val) data points were visualized in Volcano plots generated by Prism 9. For the SMC1 technical replicate experiment, a p-value <0.01 cut off from for significant hits was applied while a traditional p-value <0.05 cut off for significant hits was used for the SMC2 and HUVEC process replicate experiments. Initial filtering of significant hits from both the adj.P.Val.mWT.over.mEQ and adj.P.Val.mWT.over.Luc identified regulated peptides associated with ADAMTS7 protease activity. No logFC cut-off was applied, but positive logFC.mWT.over.mEQ values predicted candidate cleavage sites. In some cases, the same substrate cleavage site was identified from multiple peptides representing missed cleaved and fully cleaved at the C-termini versions of the peptide. To collapse the discovery set in to a single unique site, the geneSymbol field was concatenated with the StartAA field to generate a cleavage site identifier. Additional filtration of ADAMTS7 auto-catalytic sites with removal of significant hits from adj.P.Val.mEQ.over.Luc (associated with overexpression of the catalytically inactive mutant) was performed to generate a high confidence discovery set for each TAILS experiment. Overlap analysis between the SMC1, SMC2 and HUVEC TAILS experiments was performed to identify consistently regulated peptides as candidate ADAMTS7 substrate cleavage sites. Venn diagrams were made manually in Adobe Illustrator. Analysis of cleavage site positions -4 to +4 was performed with Weblogo (31) and iceLogo (32) to generate logo consensus and cleavage site heat maps factoring in the natural abundance of each amino acid in the human proteome.

### EFEMP1 Substrate Validation Experiments

Cultured HUVEC with high endogenous EFEMP1 (Fibulin-3) expression were chosen for cell-based validation experiments of TAILS identified cleavage sites. Media from HUVEC transduced with 50 MOI Ad-Luc, Ad-mADAMTS7 WT or Ad-mADAMTS7 EQ was collected in serum free conditions (Lifeline Basal EC +bFGF2 10ng/ml) and concentrated using 3kDa Amicon spin columns. Ten percent of the concentrated media was run on a 4-20% Mini-PROTEAN gel (Bio-Rad) and analyzed by western blot to detect the carboxyl region of EFEMP1/Fibulin-3 (antigen 140-209, Novus, NBP2-57871) or ADAMTS7-Flag using M2-HRP. Remaining concentrated media was run on a parallel gel for Coomassie blue staining to collect size specific bands. Gel slices were digested with trypsin or chymotrypsin for mass spectrometry analysis of semi-trypsin and semi-chymotrypsin peptides (Whitehead Institute Proteomics Core Facility). Unique EFEMP1 non-tryptic or non-chymotryptic sites identified more than once from the combined gel slices were compared across Luc, WT and EQ samples. The number of unique mass spectrometry identified peptides representing a potential cleavage site and the combined area for these peptides was used as a semi-quantitative method to identify and compare the proportion of EFEMP1 123.124 and EFEMP1 124.125 cleavage sites. To validate EFEMP1 cleavage in a binary assay, purified recombinant human HA-tagged EFEMP1/Fibulin-3 (R&D, 8416-FB) provided in PBS was dialyzed into TBS pH 8.0 to prevent precipitation with the CaCl2 in the assay buffer. Purified mouse Adamts7 WT S3A 3xFlag contained a 250 kDa full-length form and 150 kDa truncated Flag tagged form enriched in later SEC fractions as previously described (28). Purified mouse ADAMTS7 S3A proteins (0.5ug) were incubated with HA-EFEMP1/Fibulin-3 (1.0ug) at 37C in an assay buffer containing 50mM Tris pH 8.0, 150mM NaCl, 5mM CaCl2, 10uM ZnCl2 and 0.004% Bridj35. The carboxyl region of EFEMP1/Fibulin-3 was detected with Novus antibody NBP-57581 and the amino terminal HA-tag was detected with anti-HA antibody (Cell Signaling, C29F4). Coomassie stained gel slices were submitted for mass spectrometry analysis of semi-trypsin and semi-chymotrypsin peptides (Whitehead Institute Proteomics Core Facility). Analysis was performed similar to the HUVEC validation experiment to identify and compare the proportion of EFEMP1 1223.124 and EFEMP1 124.125 cleavage sites.

## Results

### ADAMTS7 TAILS Experimental design

All TAILS experiments were constructed to allow for comparison of the active WT protease in triplicate with either of the two negative control groups (supplemental Fig. 1). Control groups contained either the expression of a non-specific luciferase or an inactive EQ catalytic mutant protease with a glutamate to glutamine substitution in the H<u>E</u>xxH metalloproteinase active site (28). We used three different experimental designs to perform TAILS discovery experiments in human coronary artery smooth muscle cells (SMC) and human umbilical vein endothelial cells (HUVEC) (supplemental Fig. 1*A-D*). In the first experiment, referred to as SMC1, media was pooled from one of three different conditions (Ad-Luc control, Ad-mADAMTS7 WT or Ad-mADAMTS7 EQ) and then split into three technical replicates for each condition to minimize biological variation. In the second experiment, referred to as SMC2, input media from each triplicate condition was processed separately. In the third experiment, process replicates from adenovirus transduced HUVEC were used to identify ADAMTS7 substrate cleavage sites from secreted factors and extracellular matrix proteins originating from vascular endothelial cells (supplemental Fig. 1*E-F*). For all three experiments the total proteome of the secretome was also measured by quantitative proteomics.

**FIG. 1.**
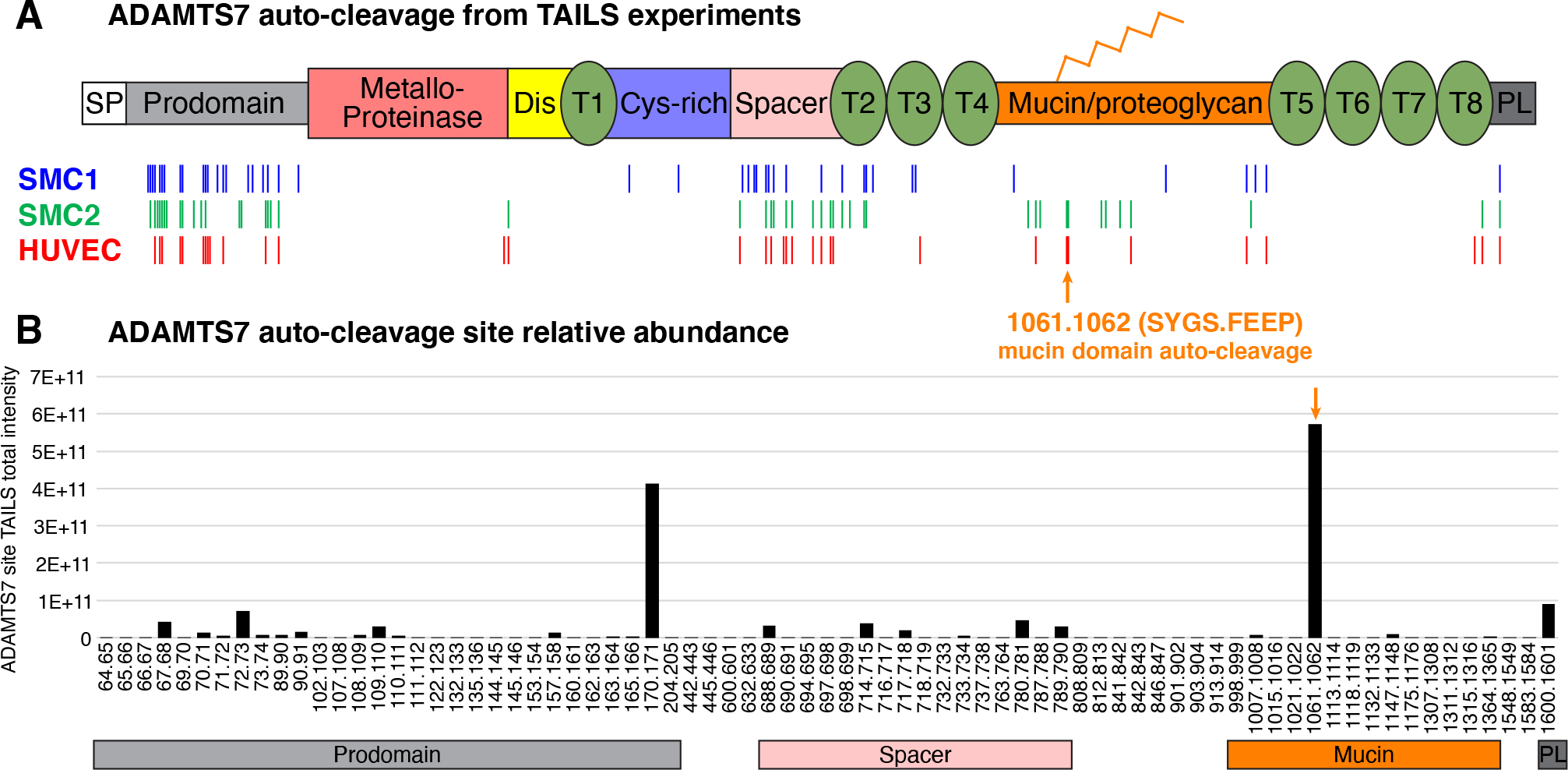
ADAMTS7 auto-cleavage sites detected in TAILS experiments. *A*, ADAMTS7 protein domains and locations of the TAILS significantly regulated WT/EQ peptides indicating auto-cleavage events. Abbreviated ADAMTS7 domains: signal peptide (SP), disintegrin (Dis), thrombospondin repeats (T), cysteine-rich (Cys-rich), protease and lacunin (PL). *B*, ADAMTS7 auto-cleavage peptide total intensities pooled from all TAILS experiments to show relative abundance of each event.

Greater than 80% of the postTAILS identified spectra contained an isobaric label to enable quantification and allowed for quantitative comparison of 8,818 peptides from 3,152 proteins, 10,964 peptides from 3,579 proteins and 13,276 peptides from 3,826 proteins in SMC1, SMC2 and HUVEC experiments, respectively (supplementary Table S1). Total secretome proteome analysis resulted in 1847 in SMC1, 1808 in SMC2 and 2031 in HUVEC fully quantitated proteins.

**Table 1.** ADAMTS7 candidate substrates identified in all TAILS experiments. High confidence ADAMTS7 cleavage sites consistently detected in all TAILS discovery datasets. Fold change enrichment with ADAMTS7 activity (logFC WT/EQ) and TAILS peptide ID total intensity displaying relative abundance in each TAILS experiment is shown for the 24 candidate sites. Sequence context for each cleavage site is shown (P4 through P4’) and location within the target candidate protein.

| TAILS_id | SMC1<br>logFC<br>WT/EQ | SMC1<br>total<br>Intensity | SMC2<br>logFC<br>WT/EQ | SMC2<br>total<br>Intensity | HUVEC<br>logFC<br>WT/EQ | HUVEC<br>total<br>Intensity | Cleavage<br>site | Substrate location |
| --- | --- | --- | --- | --- | --- | --- | --- | --- |
| AEBP1_37 | 3.41 | 4.54E+07 | 3.48 | 8.64E+07 | 4.34 | 5.26E+06 | EIEE . FLEG | N-terminal region |
| COL18A1_1505 | 1.42 | 1.68E+09 | 1.47 | 1.11E+10 | 3.88 | 1.34E+11 | EVAA . LQPP | NC1 hinge region |
| EFEMP1_124 | 4.43 | 3.19E+09 | 3.27 | 2.58E+10 | 3.31 | 2.92E+11 | ASAA . AVAG | Within atypical EGF1 |
| EFEMP1_125 | 2.25 | 2.03E+09 | 2.42 | 7.46E+09 | 2.45 | 9.33E+10 | SAAA . VAGP | Within atypical EGF1 |
| FBN1_29 | 4.49 | 1.26E+10 | 3.57 | 2.40E+10 | 3.91 | 1.85E+09 | ADAN . LEAG | N-term/propeptide |
| FLNA_1285 | 5.06 | 5.65E+08 | 5.72 | 1.38E+09 | 3.81 | 2.92E+09 | DARA . LTQT | Filamin 11 |
| FN1_36 | 3.34 | 4.40E+10 | 3.69 | 1.54E+11 | 3.45 | 1.76E+11 | QAQQ . MVQP | N-terminal region |
| FN1_40 | 4.27 | 3.67E+09 | 2.66 | 3.29E+09 | 2.96 | 1.37E+10 | MVQP . QSPV | N-terminal region |
| FN1_1143 | 0.86 | 1.36E+08 | 2.80 | 9.55E+08 | 3.68 | 4.67E+09 | VVSG . LTPG | Fibronectin type-III 6 |
| FN1_1656 | 1.52 | 4.82E+09 | 1.86 | 2.00E+10 | 1.63 | 2.23E+10 | KWLP . SSSP | Fibronectin type-III 12 |
| HSPG2_275 | 2.49 | 2.35E+07 | 2.10 | 1.16E+08 | 0.77 | 2.67E+07 | APQP . LLPG | Linker (LDL-A1/A2) |
| HSPG2_1863 | 2.03 | 1.64E+08 | 1.90 | 1.42E+08 | 2.53 | 4.95E+08 | ASGT . LSAP | Ig-like C2-type 3 |
| HSPG2_2331 | 2.04 | 7.64E+07 | 1.91 | 2.66E+08 | 1.73 | 2.45E+09 | GTQG . ANLA | Ig-like C2-type 8 |
| IGFBP7_101 | 3.69 | 5.68E+07 | 2.57 | 4.97E+08 | 1.89 | 1.12E+08 | KAGA . AAGG | IGFBP N-term region |
| LOX_124 | 3.62 | 1.65E+08 | 3.45 | 1.64E+09 | 3.35 | 8.74E+07 | TARH . WFQA | N-term/propeptide |
| LOXL2_37 | 3.28 | 5.20E+09 | 3.30 | 9.09E+09 | 3.48 | 1.61E+09 | YPEY . FQQP | N-terminal region |
| LTBP1_393 | 3.89 | 9.30E+08 | 6.30 | 6.93E+09 | 3.33 | 9.19E+08 | AADT . LTAT | Linker (EGF1/2) |
| LTBP3_239 | 2.62 | 1.25E+08 | 1.59 | 8.70E+08 | 1.57 | 5.41E+08 | PPEA . SVQV | Linker (EGF1/TB1) |
| MAP4_815 | 1.19 | 5.58E+08 | 1.70 | 5.60E+08 | 2.51 | 6.08E+08 | TTAA . AVAS | Pro-rich linker |
| MAP4_816 | 1.62 | 1.57E+08 | 2.51 | 8.83E+08 | 2.05 | 2.73E+08 | TAAA . VAST | Pro-rich linker |
| NID2_291 | 4.04 | 2.94E+09 | 3.16 | 6.67E+09 | 2.90 | 5.96E+08 | LSAA . HSSV | Linker (NIDO/EGF1) |
| SERBP1_60 | 2.10 | 2.09E+08 | 1.62 | 2.49E+08 | 0.94 | 2.20E+08 | QAAA . QTNS | N-terminal region |
| SERBP1_116 | 2.55 | 4.56E+08 | 3.25 | 7.27E+09 | 2.44 | 4.39E+09 | RPDQ . QLQG | N-terminal region |
| SRGN_59 | 4.33 | 2.70E+09 | 3.66 | 2.86E+09 | 1.17 | 2.32E+08 | PMFE . LLPG | N-term (after disulfide) |

### Statistical analysis of the datasets

TMT reporter ion ratio of each sample to the pooled reference was used for the quantitative analysis. Data was normalized using median ratio of TMT labeled natural N-termini peptides prior to application of moderated two sample T-test for comparison of the three groups for each TAILS experiment (supplementary Table S2). As expected, correlation of the technical replicates in SMC1 experiment was higher than the process replicates in SMC2 and HUVEC experiments (supplemental Fig. S2*A-C*). Therefore, we used multiple hypothesis tested adjusted p-value (adj. p-val) threshold of 0.01 for the SMC1 experiment and 0.05 for the SMC2 and HUVEC experiments. Results of the two-sample T test are shown as volcano plots in supplemental Fig. S3*A-C*. Differentially regulated peptides in WT/EQ, WT/Luc and EQ/Luc comparisons passing the significance threshold for each experiment were predominantly in the positive log fold change (logFC) side of each TAILS volcano plot. Peptides identified as mouse ADAMTS7 are colored in red for each plot.

Significantly regulated peptides in the WT/EQ and WT/Luc comparisons with positive logFC values represent neo-N-termini potentially associated with ADAMTS7 catalytic activity. In contrast, the EQ/Luc significant and positive logFC neo-N-termini are more likely to be artifacts from adenoviral expression of a full-length catalytically inactive protein. A cluster analysis of the significantly regulated TAILS peptides (supplemental Fig. S2*D-F*) revealed that the Luc and EQ negative controls clustered together and away from the active WT condition.

Secretome proteome datasets followed the same trends as the TAILS datasets with the highest correlation of replicates observed in SMC1 experiment (supplemental Fig. S4*A-C*). Normalized TMT labeled secretomes were analyzed similar to the TAILS experiment and a moderated two sample T-test was used to compare the three groups for each total secretome experiment (supplementary Table S3). Cluster analysis of the secretome proteins showed more similarity in the secreted proteomes between the WT and EQ expressing samples than the Luc control sample (supplemental Fig. S4*D-F*). Therefore, the secretomes clustered not with ADAMTS7 activity, as with the TAILS experiments, but with ADAMTS7 expression regardless of catalytic activity. Comparative secretome analysis of the WT/EQ, WT/Luc and EQ/Luc conditions was performed using the same significance thresholds from the corresponding TAILS experiments and visualized using volcano plots (supplemental Fig. 3*D-F*). The position of mouse ADAMTS7 protein was noted in each volcano plot with a red diamond. As expected, mouse ADAMTS7 was significantly upregulated in WT/Luc and EQ/Luc comparisons, with a logFC range of 2.5-3 for the SMC experiments and a logFC range of 8-9 for the HUVEC experiment. In contrast there was no significant expression difference in the WT/EQ secretome comparisons for mouse ADAMTS7 indicating a similar level of WT and EQ protein abundance. Next, we filtered the significantly regulated proteins to logFC >1 (greater than 2-fold upregulated) or <-1 (down regulated more than 2-fold) and looked for commonalities in the WT/EQ, WT/Luc and EQ/Luc comparisons using a Venn diagram for each experiment (supplemental Fig. S5*A-C*). Using these criteria, very few proteins were associated purely with proteolytic activity from the WT/EQ secretomes. Furthermore, there were no commonalities between the three secretome experiments with the exception of mouse ADAMTS7 in the overlap between WT/Luc and EQ/Luc.

### Examination of ADAMTS7 auto-cleavage events

Next, we focused on the regulated peptides from mouse ADAMTS7 in the TAILS experiments. Of the significantly regulated peptides for the WT/EQ comparisons associated with ADAMTS7 activity from each experiment, around 17% corresponded to mouse ADAMTS7 peptides and nearly all were upregulated in WT sample (supplemental Fig. 3*A-C*). The WT/EQ ADAMTS7 peptides that were significantly enriched were also found in the WT/Luc comparisons associated with ADAMTS7 activity.

Therefore, these ADAMTS7 neo N-termini from the WT/EQ comparison were likely due to *cis* or *trans* auto-cleavage events. On the other hand, there were EQ/Luc regulated peptides that were also present in the WT/Luc comparisons, possibly representing proteolysis from other cellular proteases. Consistent with this hypothesis, the total number of WT/EQ and EQ/Luc significant ADAMTS7 logFC+ peptides added up to roughly the number of WT/Luc logFC+ peptides (supplementary Table S4). WT/EQ significantly enriched peptides filtered against EQ/Luc significant peptides resulted in our list of ADAMTS7 auto-cleavage sites from each TAILS experiment (supplementary Table S5).

We compared the locations of the prospective auto-cleavage events from each TAILS experiment and found a third to nearly half of the unique sites were located in the prodomain (residues 21-220). Only 11 of the ADAMTS7 sites were common to all experiments, with 7 in the prodomain, 3 in the spacer domain and 1 in the PLAC domain (Fig. 1*A*). The most abundant prospective auto-cleavage sites based on spectral total intensity were found at 170.171 (HAQP|HVVY) in the prodomain, 1061.1062 (SYSG|FEEP) in the mucin domain and 1600.1601 (EDCE|LVEP) in the PLAC domain (Fig. 1*B* and supplementary Table S5). It is notable that the predominant TAILS identified ADAMTS7 auto-cleavage site is identical to the mucin domain auto-cleavage product we previously reported (28) and serves to validate our approach for identifying ADAMTS7 dependent cleavage sites.

### ADAMTS7 substrate cleavage sites identified by TAILS

To generate a high confidence list of substrates from each TAILS experiment, we applied a series of requirements and filters to the significant hits from the WT/EQ comparisons (see Methods). First, we excluded any ADAMTS7 sites or any sites with a logFC less than zero from the WT/EQ significant hits (supplementary Table 4). Second, we performed the same exclusions to the WT/Luc significant hits as a separate comparison for ADAMTS7 function and used the filtered overlap from the WT/EQ and WT/Luc as a stringent constraint for ADAMTS7 catalytic activity. Lastly, we removed any sites that were significantly upregulated in the EQ/Luc comparisons or any duplicate identifications from multiple peptides to generate a unique list of high confidence substrate cleavage sites. Volcano plots with the mouse ADAMTS7 peptides removed from the dataset show the high confidence substrate cleavage site regulated peptides (labeled in green) within the upper right quadrant (Fig. 2*A-C*). Histograms illustrate the overlap of significantly regulated unique cleavage sites from each of the comparisons and display a similar trend for the independent TAILS experiments (Fig. 3*A,C,E*). It is noteworthy that the WT/EQ high confidence regulated peptides on average were more significant and had higher logFC values compared to WT/EQ only regulated peptides which lacked independent verification within each TAILS dataset (supplemental Fig. S6).

**FIG. 2.**
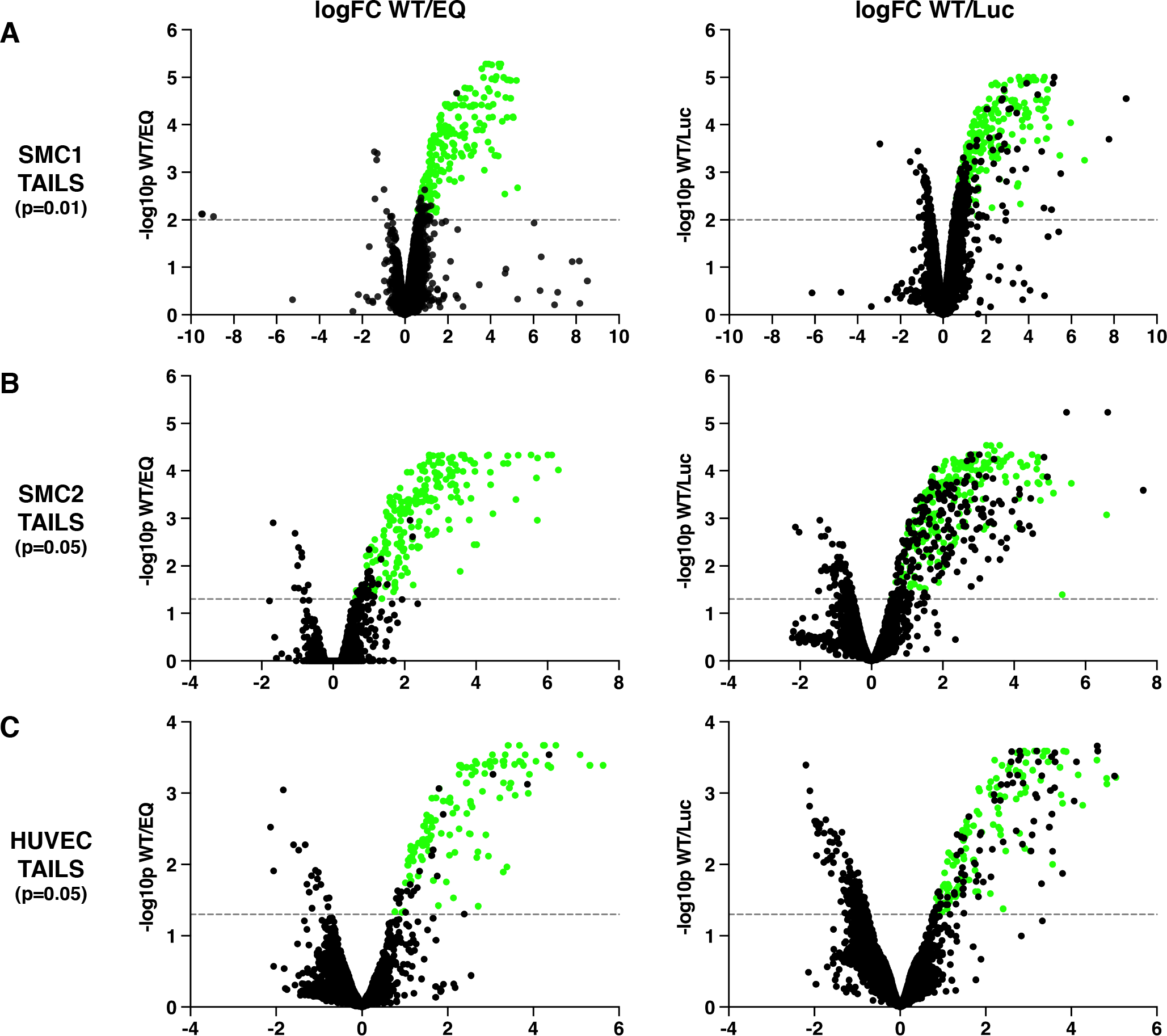
Volcano plots of TAILS regulated peptides visualizing the high confidence cleavage sites. *A-C*, comparison of WT/EQ and WT/Luc regulated peptides from three independent TAILS experiments after removal of mouse ADAMTS7 peptides. *A*, TAILS SMC1 (p<0.01) *B*, TAILS SMC2 (p<0.05) *C*, TAILS HUVEC (p<0.05), dotted line reflects significance cut off on the -log10P scale. Regulated peptides enriched for ADAMTS7 WT activity and meeting all criterial for the high confidence candidate cleavage sites (signifincant for both WT/EQ and WT/Luc comparisons) are shown in green for each TAILS experiment.

From the original number of WT/EQ significantly regulated peptides, 66% of the SMC1, 67% of the SMC2 and 60% of the HUVEC significant hits passed all these criteria (supplementary Table S4 and supplementary Table S6). Next, we compared the substrate logo and heat maps of high confidence substrate cleavage sites from the SMC and HUVEC TAILS experiments.

Based on the iceLogo plots, AA|L at the P2, P1 and P1’ positions was most commonly observed (Fig. 3*B,D,F*); however, each of these amino acids was present at no more than 30% in their respective positions (supplemental Fig. S7). Although a strict consensus was not apparent from our individual high confidence datasets, similarities were evident in candidate substrate heat maps and amino acid frequency plots from different TAILS experiments (supplemental Fig. S8).

### TAILS discovery set overlap analysis

By virtue of having multiple independent TAILS discovery experiments for ADAMTS7, we compared the list of high confidence substrate cleavage sites to emphasize those that were found more than once. From the comparison of substrate cleavage sites from SMC1 (207 identified solely by trypsin digests) and SMC2 (210 identified from half trypsin or half AspN digests), 66 were identical resulting in an overlap of 32% of the identified sites (Fig. 4*A*). Although both SMC samples were digested with trypsin, 13 of the 66 unique sites were identified by AspN regulated peptides from the SMC2 TAILS experiment rather than the comparable semi-tryptic peptide from the SMC1 TAILS experiment, demonstrating that multiple peptides can identify the same cleavage site. Sorting the overlapping SMC cleavage sites into common genes identified multiple cleavage sites in several candidate substrates, including 8 for COL1A2 (Collagen type I alpha-2 chain), 6 for FN1 (Fibronectin), 4 for HSPG2 (Basement membrane-specific heparan sulfate proteoglycan core protein/Perlecan) and 4 for LOX (Protein-lysine 6-oxidase). Comparison of the candidate sites from SMC2 and HUVEC (127, also identified from half trypsin or half AspN digests) resulted in 44 identical cleavage sites, including additional sites in FN1, HSPG2 and LOX not present in the SMC1 TAILS high confidence cleavage site list (Fig. 4*B*). Of these 44 shared sites, the 20 that were unique to the SMC2/HUVEC comparison were predominantly from the AspN digestion (15 semi-AspN peptides and 5 purely semi-tryptic peptides), which may explain why these sites were not identified in the SMC1 trypsin only digestion TAILS. In total there were 91 unique sites identified multiple times in our TAILS discovery experiments for ADAMTS7 substrates (supplementary Table S6). When the individual sites are grouped into their corresponding gene, 48 potential substrates emerged (Fig. 4*C*). The most unique cleavage sites from multiple datasets were identified from FN1, including 4 unique cleavage sites from significantly regulated peptides in all TAILS experiments. Furthermore, both FN1 and LOX are amongst the CAD risk loci categorized as non-lipid vascular remodeling pathway genes along with ADAMTS7 (33).

**FIG. 3.**
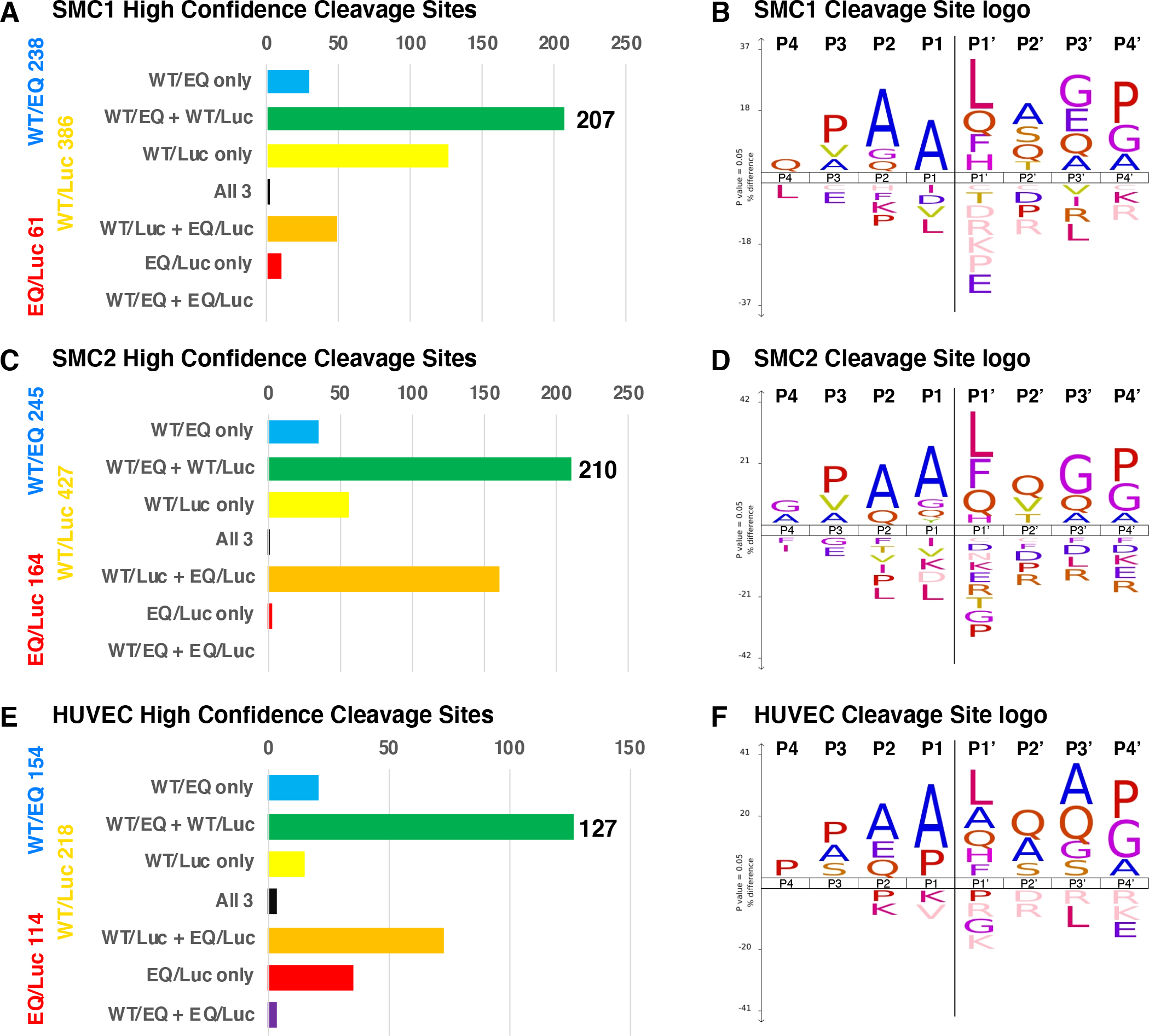
ADAMTS7 TAILS high confidence cleavage sites from independent experiments. Histograms showing the overlap between significantly regulated candidate cleavage sites from the SMC1 (*A*), SMC2 (*C*) and HUVEC (*E*) TAILS experiments. Candidate cleavage sites present in both the WT/EQ and WT/Luc comparisons were consistently associated with ADAMTS7 activity and are defined as high confidence cleavage sites (shown as green dots in Figure 2). The remaining regulated peptides were significant for one or more condition(s) in the histogram categories. Analysis of the cleavage sites using iceLogo shows the similarities between independent TAILS experiments for SMC1 (*B*), SMC2 (*D*) and HUVEC (*F*).

**FIG. 4.**
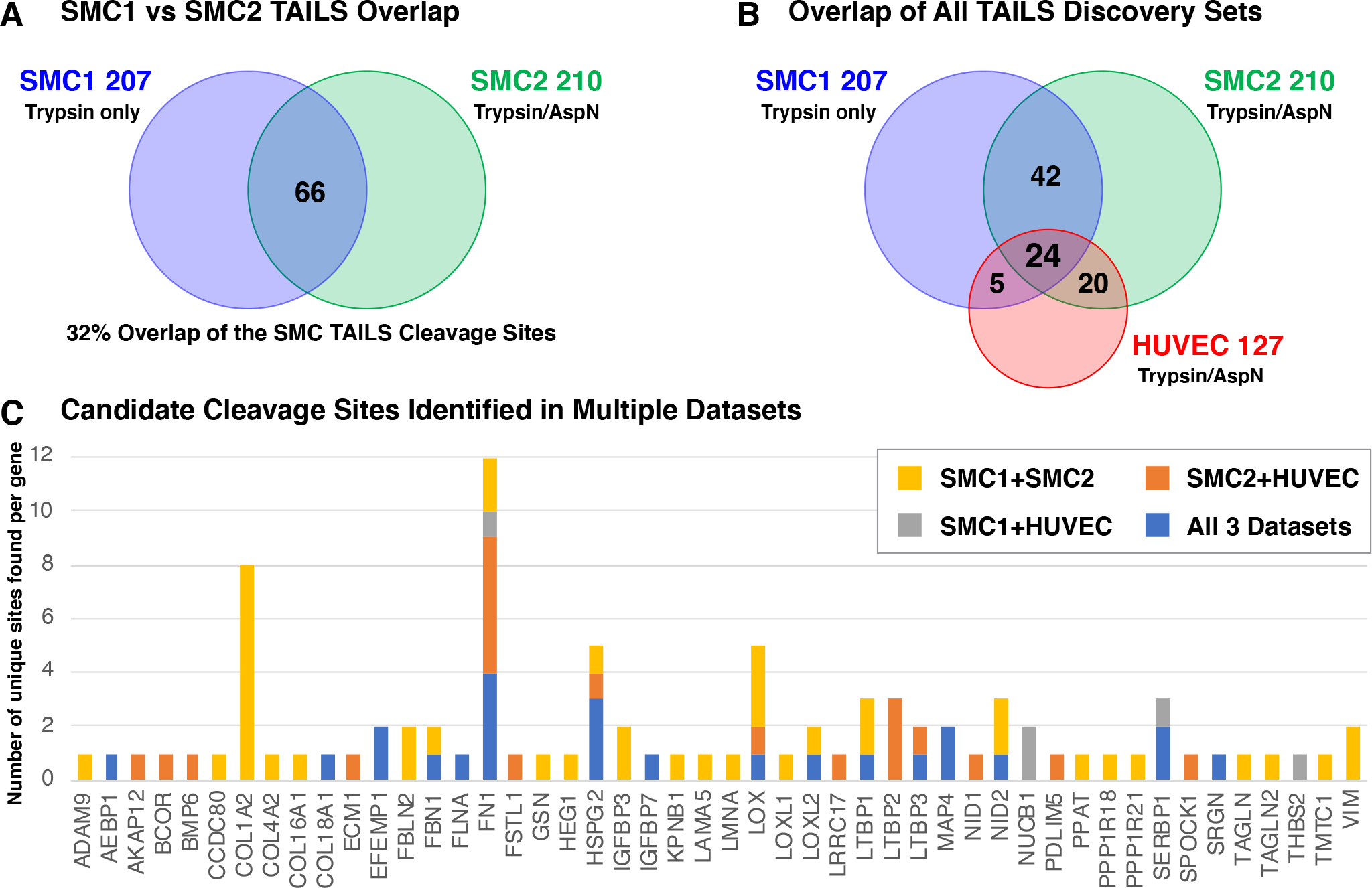
ADAMTS7 TAILS Discovery Set Overlap Analysis. *A*, Venn diagram showing the overlap of unique candidate cleavage sites from SMC1 and SMC2 high confidence sites. *B*, Venn diagram showing the overlap from all SMC and HUVEC TAILS datasets. *C*, Gene assignment of the 91 unique candidate cleavage sites identified from multiple TAILS experiments, including 24 unique sites from 16 different genes identified in all three TAILS datasets.

Analyzing the overlap between all three TAILS discovery sets identified 24 unique cleavage sites encoded by 16 different genes (Table 1). Most ADAMTS7 candidate substrates from this list were primarily localized in the extracellular region, with the exception of proteins with defined roles in the cytoskeletal (FLNA and MAP4) and nuclear (SERBP1) intracellular regions. The 24 cleavage sites were found in a variety of substrate protein domains and were commonly found in N-terminal regions or unstructured linker regions. Remarkably some of the unique cleavage sites were found at adjacent positions in the same candidate substrate, as was the case with EFEMP1 and MAP4. In both cases the logFC ratios favored the more N-terminal cleavage site, which may indicate either an initial preference for the first site or a sequential cleavage event while the enzyme remained associated with the cleaved substrate. By analyzing the overlap between independent TAILS discovery sets, we have identified reproducible substrate cleavage sites which we used to prioritize validation experiments to confirm ADAMTS7 protease specificity.

### Validation of ADAMTS7 preference for EFEMP1 cleavage at adjacent sites

EFEMP1 is a secreted extracellular matrix protein with multiple EGF domains and carboxyl terminal Fibulin domain. The first EGF domain is atypical and contains an extended linker region with documented sensitivity to proteases; and is also the location of candidate ADAMTS7 123.124 and 124.125 cleavage sites (Fig. 5*A*). By comparison, a previous TAILS experiment with ADAMTS3 reported EFEMP1 cleavage at 122.123 and 123.124, although no EFEMP1 cleavage sites were identified in TAILS experiments for paralogs ADAMTS2 or ADAMTS14 (25). MMP3 and MMP7 are reported to cleave EFEMP1 at 124.125, and notably there was an observed background cleavage at 123.124 within experiments from the same study (34). Data from our TAILS experiments consistently showed a higher logFC and total intensities for the 123.124 site, predicting 2-3 fold more cleavage at 123.124 (ASAA|AVAG) compared to the adjacent 124.125 (SAAA|VAGP) site (Table 1). To verify that EFEMP1 is a substrate for ADAMTS7 and to compare the relative amount of cleavage at the 123.124 and 124.125 sites, we performed two key validation experiments.

**FIG. 5.**
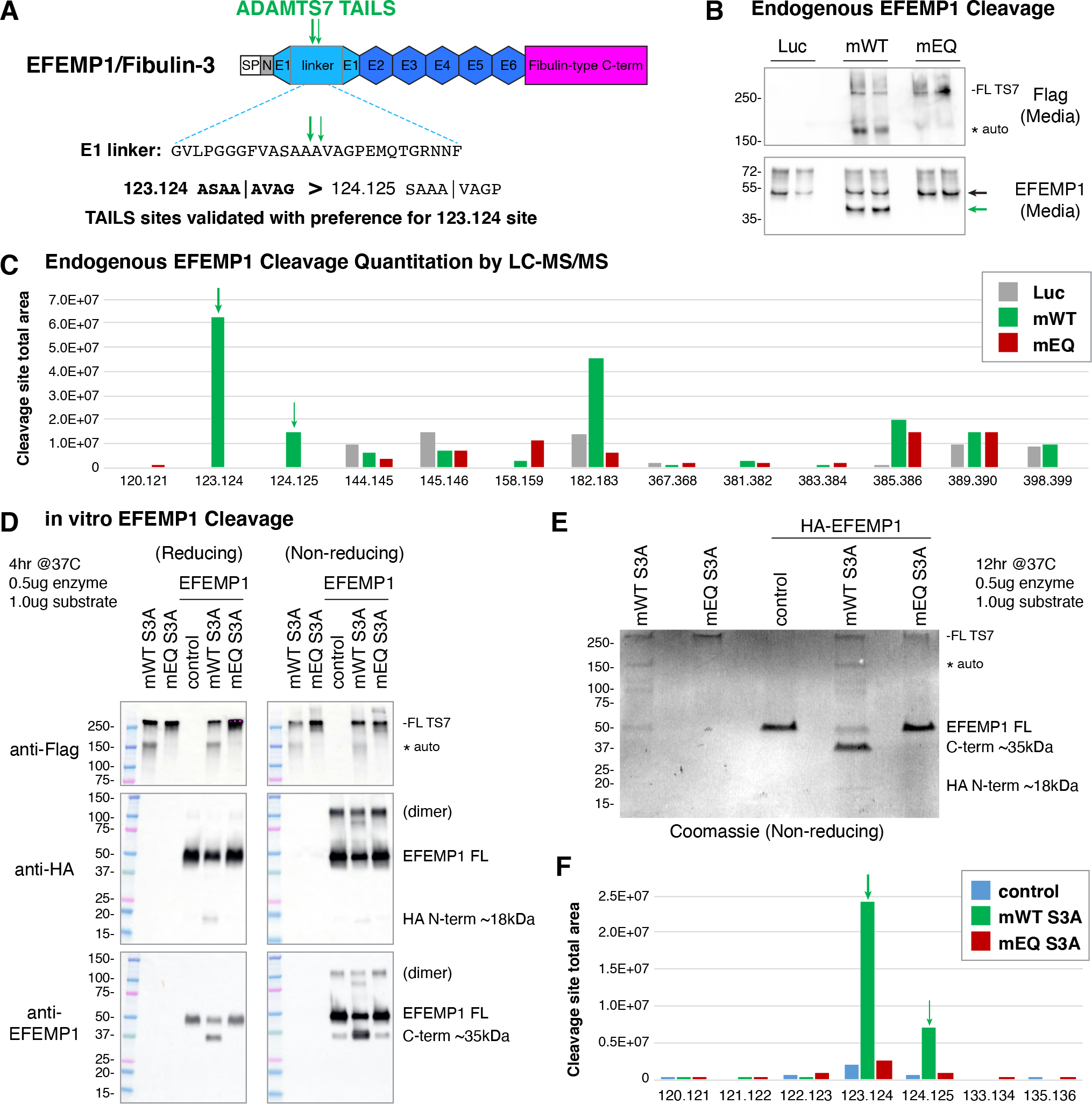
Validation of TAILS substrate EFEMP1 and cleavage site preference. *A*, EFEMP1/Fibulin-3 protein domains, amino acid sequence of the atypical EGF repeat linker and location of the ADAMTS7 cleavage sites. Abbreviated EFEMP1 domains: signal peptide (SP), N-terminal region (N), EGF repeats (E). *B*, concentrated media from HUVEC expressing Ad-Luc, Ad-mWT or Ad-mEQ assessed by western blot under non-reducing conditions. Anti-EFEMP1 antibody recognizes an epitope C-terminal to the ADAMTS7 cleavage sites. *C*, quantitation of semi-tryptic or semi-chymotryptic peptides from HUVEC media matching novel cleavage sites from the endogenous EFEMP1 protein. The total area was greater for the 123.124 cleavage site compared to the adjacent 124.125 cleavage site. Additional cleavage events observed were also found in the Luc and EQ controls. *D*, in vitro cleavage of HA-EFEMP1 by purified full-length mouse ADAMTS7 S3A assessed by western blot. The antibodies to the N-terminal HA epitope and C-terminal EFEMP1 epitope recognized the EFEMP1 more strongly under non-reducing conditions. A band at 100kDa under non-reducing conditions is consistent with a purified HA-EFEMP1 dimer, which was also sensitive to ADAMTS7 cleavage. *E*, overnight digest of HA-EFEMP1 by mouse ADAMTS7 S3A assessed by Coomassie staining. *F*, quantitation of semi-tryptic or semi-chymotryptic peptides from the atypical EGF1 repeat region from HA-EFEMP1 showing a consistent preference for the 123.124 cleavage site.

We first examined the specificity of ADAMTS7 cleavage to endogenously expressed EFEMP1. HUVEC express higher levels of EFEMP1 compared to SMC, as shown by higher total intensities of EFEMP1 spectra from the secretome experiments (supplemental Table S3). Similar to the initial HUVEC discovery experiment, we expressed full-length mouse ADAMTS7 WT and EQ using adenovirus and examined the concentrated media. As expected ADAMTS7 WT migrated at two bands matching the observed autocleavage event in the mucin domain (Fig. 5*B*). Using an antibody to an EFEMP1 epitope c-terminal to the candidate ADAMTS7 cleavage sites, we detected a single lower mobility band consistent with cleavage within the atypical first EGF domain. MS analysis of these bands following digestion with trypsin or chymotrypsin identified semi-tryptic and semi-chymotryptic peptides indicating cleavage at 123.124 and 124.125 sites from the Ad-ADAMTS7 WT sample. Importantly, these sites were not detected in the Luc or EQ samples while our methods were sensitive enough to identify cleavage outside of the first EGF repeat in all samples which were presumably independent of ADAMTS7 activity. Using the area for each of these observed spectra cumulatively, we assigned a relative abundance to each of these detected cleavage products and observed a preference for 123.124 over 124.125 (Fig. 5*C*).

Next, we analyzed the site and preference of EFEMP1 cleavage by ADAMTS7 in a binary *in vitro* system. We obtained commercially purified full-length epitope tagged HA-EFEMP1 and combined it with our purified full-length mouse ADAMTS7 S3A WT or ADAMTS7 S3A EQ for 4 hours at 37C. Mobility of EFEMP1 was detected by western blot using samples in reducing or non-reducing conditions. Full-length HA-EFEMP1 was detected at the expected position and a lower band was detected matching the predicted HA tagged amino terminus after cleavage by ADAMTS7 in the atypical first EGF domain (Fig. 5*D*). The antibody to the EFEMP1 carboxyl terminus provided a stronger signal under non-reducing compared to reducing conditions. Here we could detect an abundant lower band at 40kDa consistent with ADAMTS7 cleavage under reducing and non-reducing conditions. A faint band at this position was also present in the no enzyme and ADAMTS7 EQ control reactions under non-reducing conditions raising the possibility of background cleavage from the commercial EFEMP1. We performed a second experiment with digestion for 12 hours and examined the products by Coomassie staining (Fig. 5*E*). In the WT treated condition we could detect two lower bands matching to the amino and carboxyl terminal fragments of EFEMP1, consistent with ADAMTS7 cleavage restricted to the atypical first EGF domain. Corresponding bands from this gel were analyzed by mass spectrometry to investigate the relative abundance of these events. Here we identified the suspected background cleavage under conditions with no enzyme or with ADAMTS7 S3A EQ treatment, however these occurrences were relatively minor compared to the activity stimulated by ADAMTS7 S3A WT (Fig. 5*F* and supplementary Fig. S9). From this binary *in vitro* system, we were able to demonstrate ADAMTS7 mediated cleavage of EFEMP1 in the atypical first EGF domain and confirm a preference for 123.124 over 124.125 cleavage.

In total, our detection and quantitation of both endogenous and purified EFEMP1 are consistent with the findings from three independent TAILS discovery experiments to identify new ADAMTS7 substrates.

## Discussion

In this study we performed TMT-TAILS to identify substrates for ADAMTS7 from the secretomes of vascular smooth muscle and endothelial cells. Each of the three independent TAILS experiments identified previously unknown candidate substrate cleavage sites associated with ADAMTS7 activity. To confirm our findings, we presented the validation of two adjacent cleavage sites 123.124 (ASAA|AVAG) and 124.125 (SAAA|VAGP) within the atypical first EGF repeat of EFEMP1.

EFEMP1, commonly known as Fibulin-3, is a secreted extracellular matrix protein highly expressed in the vasculature in a pattern overlapping with ADAMTS7 (GTEx Portal V8).

Targeted mutation of mouse *Efemp1* resulted in a viable knockout mouse with abnormal connective tissue due to impaired elastogenesis, including a propensity for hernias and early aging phenotypes (35). Similar connective tissue disorders were found in a human patient with *EFEMP1* truncating mutations (36). In contrast, a recurrent R345W gain-of-function mutation in the central region of *EFEMP1* results autosomal dominant Doyne honeycomb retinal dystrophy (37). Thus, there are no readily apparent correlations between the known *EFEMP1* gain- or loss-of-function phenotypes and the atheroprotection conferred from *ADAMTS7* loss-of-function. While *Adamts7* knockout mice do not have a reported connective tissue disorder, a recent report describes abnormal collagen fibrillogenesis in *Adamts7/Adamts12* double knockout mice resulting in tendon heterotopic ossification (38). Compensation and substrate redundancies between the paralogs ADAMTS7 and ADAMTS12 may explain the lack of overlap from the individual enzyme loss-of-function phenotypes and the diseases associated with mutations in the candidate substrates.

Although it is presently unclear if ADAMTS7 regulated EFEMP1 cleavage will impact vascular phenotypes, experimental evidence shows that EFEMP1 124.125 cleavage by MMP may alter interacting binding partners (34). Within the family of fibulin proteins, EFEMP1/Fibulin-3 is similar in structure to EFEMP2/Fibulin-4 and FBLN5/Fibulin-5 (37), however neither Fibulin-4 or Fibulin-5 were identified as ADAMTS7 candidate substrates from our experiments. In the related family member FBLN2/Fibulin-2, adjacent sites 258.259 (TAAA|ALGP) and 259.260 (AAAA|LGPP) in the N-terminal cysteine-free region were identified as candidate sites in both of the SMC TAILS experiments. Cleavage by ADAMTS7 at this location would release the FBLN2 N-terminal cysteine-rich domain with reported roles in FBLN2 oligomerization (39). Further work will be needed to understand the consequence of ADAMTS7 cleavage of the N-terminal regions of Fibulins 2 and 3 in disease biology.

In the active form of full-length mouse ADAMTS7, we consistently observed a lower band at 150 kDa in the media from SMC, HUVEC and HEK293 fibroblasts. Amino terminal sequencing identified the WT 150 kDa band beginning at phenylalanine 1062 (FEEPHPDL) (28). In this study, TAILS experiments digested with AspN identified significantly regulated peptides in the WT/EQ comparison to support a predominant ADAMTS7 auto-cleavage event at 1061.1062 (SYGS|FEEP) nearby the CS-GAG attachment site in the mucin domain. Removal of the amino acids 1062-1657 would preserve a CS-GAG tethered enzyme that lacks a carboxyl terminus normally thought to be required for substrate recognition, potentially changing the exosite substrate specificity for ADAMTS7.

The mouse ADAMTS7 auto-cleavage site is adjacent to one of the few highly conserved regions within the mucin domain and is partially conserved in human ADAMTS7 1080.1081 (SYGP|SEEP), although we were unable to confirm auto-cleavage for WT human ADAMTS7. Cleavage at this position was not identified in a TAILS experiment using a human ADAMTS7 truncated after the mucin domain, lacking the carboxyl terminal TSR repeats 5-8 and PLAC domains (27). From this study, they identified auto-cleavage events in the spacer domain for human ADAMTS7 (729.730 RIQE|VAEA and 732.733 EVAE|AANF) with confirmed amino terminal sequencing of the latter site. Within our TAILS datasets using mouse ADAMTS7, we identified analogous auto-cleavage sites in the spacer domain at 714.715 (LIEE|VAEA) for all TAILS experiments and 717.718 (EVAE|AANF) in the SMC2 and HUVEC TAILS experiments (supplemental Table S5). Analysis of the WT/EQ significantly enriched peptides revealed multiple sites in the prodomain consistent with auto-cleavage events, with the most abundant site at 170.171 (HAQP|HVVY). Although this site is entirely conserved in human ADAMTS7, it was not identified in the previous TAILS study (27). The ADAMTS7 prodomain contains a cysteine switch motif which acts to maintain enzyme latency through interactions with the Zinc metal in the active site (40). The mouse ADAMTS7 prodomain is processed by Furin protease at 60.61 and 220.221, with only the second Furin processing site removing the inhibitory cysteine switch at Cys194 (17). Although mouse ADAMTS7 cleavage at 170.171 would retain the cysteine switch to the catalytic domain, it is possible this may affect the progressive zymogen processing by Furin. Further work will be needed to understand how the secreted ADAMTS7 maintains latency and if auto-cleavage plays a role independent of, or in concert with, Furin processing in the prodomain.

Analysis of P4 through P4’ positions from ADAMTS7 auto-cleavage and candidate substrate cleavage sites from the TAILS experiments suggests that ADAMTS7 is able to process substrates in a variety of contexts (Fig. 3). Visually from the iceLogo plots, a preference for Alanine in the P1 position and Leucine in the P1’ was the most common, however many candidate cleavage sites identified in multiple TAILS experiments did not conform to the consensus logo plots. ADAMTS7 TAILS substrate specificity was notably similar to those reported for MMP2 TAILS experiments showing a preference for PAA|L in the P3 through P1’ positions without an absolute requirement for any of these residues (24). Compared to the preferences from other ADAMTS family members, there appear to be more similarities with the procollagen N-proteinases ADAMTS2/3/14 preference for A/P|Q (25) than the aggrecanases ADAMTS4/5 preference for E|A/L (41). In peptide based enzymatic assays, cleavage of aggrecan (TEGE|ARGS or TAQE|AGEG) or versican (EAAE|ARRG) has been reported for ADAMTS1, ADAMTS4, ADAMTS5, ADAMTS8, ADAMTS9 and ADAMTS20 (42-48). Although ADAMTS7 activity generated many candidate E|A cleavage sites within our TAILS datasets, we did not detect significantly regulated peptides for aggrecan or versican at these sites to suggest similar cleavage events. This is consistent with the initial characterization of ADAMTS7 reporting no activity for aggrecan or versican using neo-epitope antibodies for the common aggrecanase cleavage sites (17). In the previous ADAMTS7 TAILS study, a P1 glutamate preference was observed in the latent-transforming growth factor beta-binding proteins 3 and 4, specifically at LTBP3 220.221 (ISAE|VQAP) and LTPBP4 229.230 (HPQE|ASVV) (27). In our study, the LTPBP4 229.230 site was not significantly regulated in the TAILS datasets, but nearby LTBP4 233.234 (ASVV|VHQV) was significantly regulated in the SMC1 dataset. The previously reported LTBP3 220.221 site was significant in the HUVEC TAILS experiment but not in the SMC TAILS datasets. Within this region, LTPBP3 238.239 (PPEA|SVQV) was identified in all three TAILS experiments and alignment of LTBP3 and LTBP4 brings this cleavage site very close to the reported LTBP4 229.230 (HPQE|ASVV) site. Collectively, these examples from our TAILS experiments demonstrate that ADAMTS7 exhibits broad specificity for the amino acids at the site of cleavage. Nonetheless, the repeated discovery of the same diverse cleavage sites suggests that substrate specificity remains a feature of ADAMTS7 and likely relies on other non-enzymatic domains for substrate recognition.

Within our list of candidate substrate cleavage sites identified in more than one TAILS experiment, we observed that while some candidate substrates displayed cleavage within a specific region of the protein, others fell into a different category where multiple identified cleavage sites were present throughout the protein in multiple domains. This was the case for Fibronectin and HSPG2/Perlecan for all the TAILS experiments and for COL1A2 from the SMC TAILS experiments. One possibility is that ADAMTS7 normally associates with these proteins in the extracellular matrix and under our over-expression conditions opportunistically cleaves these substrates in a less regulated manner. In contrast, other identified substrates displayed a more restricted pattern of cleavage confined to a particular region which may suggest a more regulated interaction and cleavage process. This appears to be the case with EFEMP1 and may hold true for other candidate substrates such as LOX which displayed a string of candidate sites in the prodomain at positions 122.123, 123.124, 124.125 and 125.126 from multiple TAILS experiments. The candidate site that was present in all three TAILS experiments was at LOX 123.124 (TARH|WFQA) upstream from the zymogen processing site at 162.163 by the procollagen C-proteinase BMP1. The ADAMTS7 mediated prodomain LOX cleavage sites are distinct in location from those identified from the ADAMTS2/3/14 TAILS experiments or from the reported LOX catalytic domain cleavage by ADAMTS2/14 (25, 49). The LOX prodomain is essential for secretion and assists with substrate interaction. Following BMP1 cleavage, the LOX prodomain also has the ability to act as a bioactive product with tumor suppressor function independent from LOX enzymatic domain (50). Therefore, ADAMTS7 cleavage of the LOX prodomain may impact multiple functions of the pro-LOX zymogen association with substrates, pro-LOX zymogen processing or the modification/inactivation of the bioactive free LOX propeptide. In the cases of EFEMP1 and LOX, the ADAMTS7 cleavage events are at adjacent amino acid positions which may produce similar biological effects, however these phenomena may complicate the development of neo-epitope specific antibodies to a single defined cleavage site similar to the reagents developed for the aggrecanases ADAMTS4 and ADAMTS5.

In addition to inactivating structural and bioactive ECM factors, ADAMTS7 TAILS candidate cleavage sites have the potential to produce known bioactive products from unique cleavage sites in the hinge regions of COL18A1 (Collagen type XVIII alpha-1) and CTGF (Connective Tissue Growth Factor). Endostatin and endostatin-like fragments with anti-angiogenic properties originate from the carboxyl terminal region of COL18A1 following MMP/elastase/cathepsin cleavage within the hinge region (amino acids 1502-1571). The ADAMTS7 candidate cleavage site COL18A1 1504.1505 (EVAA|LQPP) found in all three TAILS experiments is located near the beginning of the hinge region, upstream from the first known MMP cleavage site at 1511.1512 by MMP7 (51). Our findings suggest that ADAMTS7 is capable of producing an endostatin-like fragment with similar anti-angiogenic activities. Additional significantly upregulated peptides corresponding to nearby cleavage at 1501.1502 (TDNE|VAAL) and 1503.1504 (NEVA|ALQP) were present in the HUVEC dataset and may correlate with the increase in COL18A1 protein levels detected in the HUVEC secretome. In fact, COL18A1 was one of the few examples a protein significantly upregulated in the WT secretome, but not in the Luc or EQ secretomes, consistent with COL18A1 upregulation in response to ADAMTS7 catalytic activity in HUVEC. This may represent a feed forward circuit with ADAMTS7 activity stimulating both the upregulation and cleavage of COL18A1 resulting in an endostatin-like matrikine.

CTGF, also known as CCN2, is a secreted multidomain matricellular protein with a central proteolytically sensitive hinge region (amino acids 168-197). It has been previously shown that cleavage in the hinge domain at 180.181 (PALA|AYRL) by MMPs generates a bioactive carboxyl terminal fragment containing the TSR and cysteine rich CT domains (52). CTGF is highly expressed in the HUVEC cell line and specifically in the HUVEC TAILS experiment we identified significantly regulated peptides representing CTGF hinge region cleavage sites at 172.173 (PKDQ|TVVG), 173.174 (KDQT|VVGP), 177.178 (VVGP|ALAA) and 186.187 (RLED|TFGP). Although multiple protease cleavage sites have been reported for the CTGF hinge region (53-55), very few known sites overlap with the ADAMTS7 TAILS candidates, with the exception of the 186.187 site identified in an ADAMTS3 TAILS study (25). CTGF was previously identified as a potential substrate for ADAMTS7 through a yeast two hybrid screen demonstrating a requirement for the ADAMTS7 mucin, TSR5-8 and PLAC domains for interaction with the CTGF amino terminal region (56). Furthermore, it was shown in an *in vitro* cleavage assay that the ADAMTS7 catalytic domain processed CTGF, producing bands compatible with cleavage in the hinge region. A similar binding interaction between CTGF and the paralog ADAMTS12 was mapped to the mucin and TSR5-8 regions of ADAMTS12, along with evidence of CTGF processing from co-transfected cells (57). Our ADAMTS7 TAILS study provides further evidence for a connection between ADAMTS7 and CTGF from an unbiased proteomic method. Based on the data from our HUVEC TAILS experiment, a cleavage site preference of 172.173 or 186.187 in the CTGF hinge region would be predicted based on logFC values and total spectra intensities. Further work will be needed to validate these, and other candidates identified from our TAILS discovery sets.

Although ADAMTS7 is characterized as a COMP protease, we were unable to identify significantly regulated peptides consistent an ADAMTS7 candidate cleavage site. COMP protein and peptides were identified in each of the TAILS and secretome experiments, however the total peptide coverage ranged from 18-23% indicating that a significant portion of COMP was not captured and quantitated in our TAILS experiments. In the case of COMP, this may be due to a limitation in the expression level of the substrate of interest in our cell lines and the depth of amino acid coverage in our experiments. Another reported ADAMTS7 substrate is thrombospondin 1 (10) and both the SMC and HUVEC cell lines expressed much higher levels of this protein resulting in 71-78% total protein coverage. Within THBS1, two regulated peptides were identified in separate experiments (629.630 in HUVEC and 971.972 in SMC1), however these were not consistently identified in the other TAILS datasets. Although our coverage for THBS1 was much higher than that for COMP, we cannot definitively rule out the possibility that a true ADAMTS7 cleavage site was missed in our TAILS experiments.

Achieving appropriate coverage and depth for a given substrate is a challenge for any unbiased degradomics technique and we attempted improve these traits in our successive TAILS experiments. For instance, in the first TAILS experiment, we digested the TMT labeled peptides with only trypsin, limiting the identification of candidate sites to peptides greater than five residues proceeding a tryptic R. or K. site that could be identified through LC-MS/MS. To improve the peptide coverage for ADAMTS7 substrates in the following TAILS experiments, we analyzed both trypsin and AspN digested products. Additionally we analyzed the peptides with a relaxed AspN condition to capture both .D or .E cleavage events to increase the number of identifiable spectra. Compared to an analysis of our datasets with a strict AspN (.D only) consensus, application of relaxed AspN condition further increased the number of quantified TMT labeled peptides by 23% for the SMC2 and 17% for the HUVEC TAILS experiments.

Including additional enzymes such as chymotrypsin with cleavage sites distinct from trypsin and AspN would likely increase depth and coverage even further to capture additional ADAMTS7 candidate cleavage sites.

Based on the diverse cleavage site specificity from ADAMTS7 in our TAILS experiments, our experimental design likely benefited from using the full-length protein with the carboxyl-terminal substrate interaction domains. Additional ADAMTS7 site specificity could be investigated using a PICS (Proteomic Identification of protease Cleavage Sites) based strategy utilizing a library of short peptides predigested with a specific enzyme like trypsin or LysC (41), however this may not reliably identify endogenous substrate cleavage sites driven by exosite specificity. Consistent with our observations of ADAMTS7 cleavage site specificity, a broad specificity for the ADAMTS7 enzyme was observed in a library of internally quenched fluorogenic peptides where nearly half were appreciably cleaved (58).

In closing, this study represents the most comprehensive list of ADAMTS7 candidate substrates from the secreted factors of vascular smooth muscle and endothelial cells. Our method using 10 isobaric tags for a single TAILS experiment shows the power of performing an individual condition in triplicate and subtractive methods to select regulated peptides consistently associated with protease activity. And finally, we show the virtue of performing multiple TAILS experiments to prioritize candidate validation experiments from the list of overlapping discovery sets.

## Acknowledgements

This work was supported by a research grant from Bayer AG within the Cardiovascular Disease Initiative at the Broad Institute. Dr. Ellinor is supported by the National Institutes of Health (1RO1HL092577, K24HL105780) and the American Heart Association (18SFRN34110082). We thank Eric Spooner at the Whitehead Institute Proteomics Core Facility for assistance with mass spec protein identification and Michael Berne at Tufts University Core Facility for assistance with protein sequencing.

## Data Availability

Mass spectrometry data from this study has been submitted to Proteome-Xchange / MassIVE. This article contains supplemental data.

**Table S1** – TAILS experiment quantitation statistics

**Table S2 –** TAILS datasets for the SMC1, SMC2 and HUVEC experiments

**Table S3 –** Secretome datasets for the SMC1, SMC2 and HUVEC experiments Table S4 – TAILS regulated peptide summary

**Table S5** – Annotated ADAMTS7 auto-cleavage sites from each TAILS experiment

**Table S6** –TAILS high confidence candidate substrate cleavage sites for each experiment and annotated overlap analysis of cleavage sites identified in multiple TAILS experiments

---

**Figure S1.**
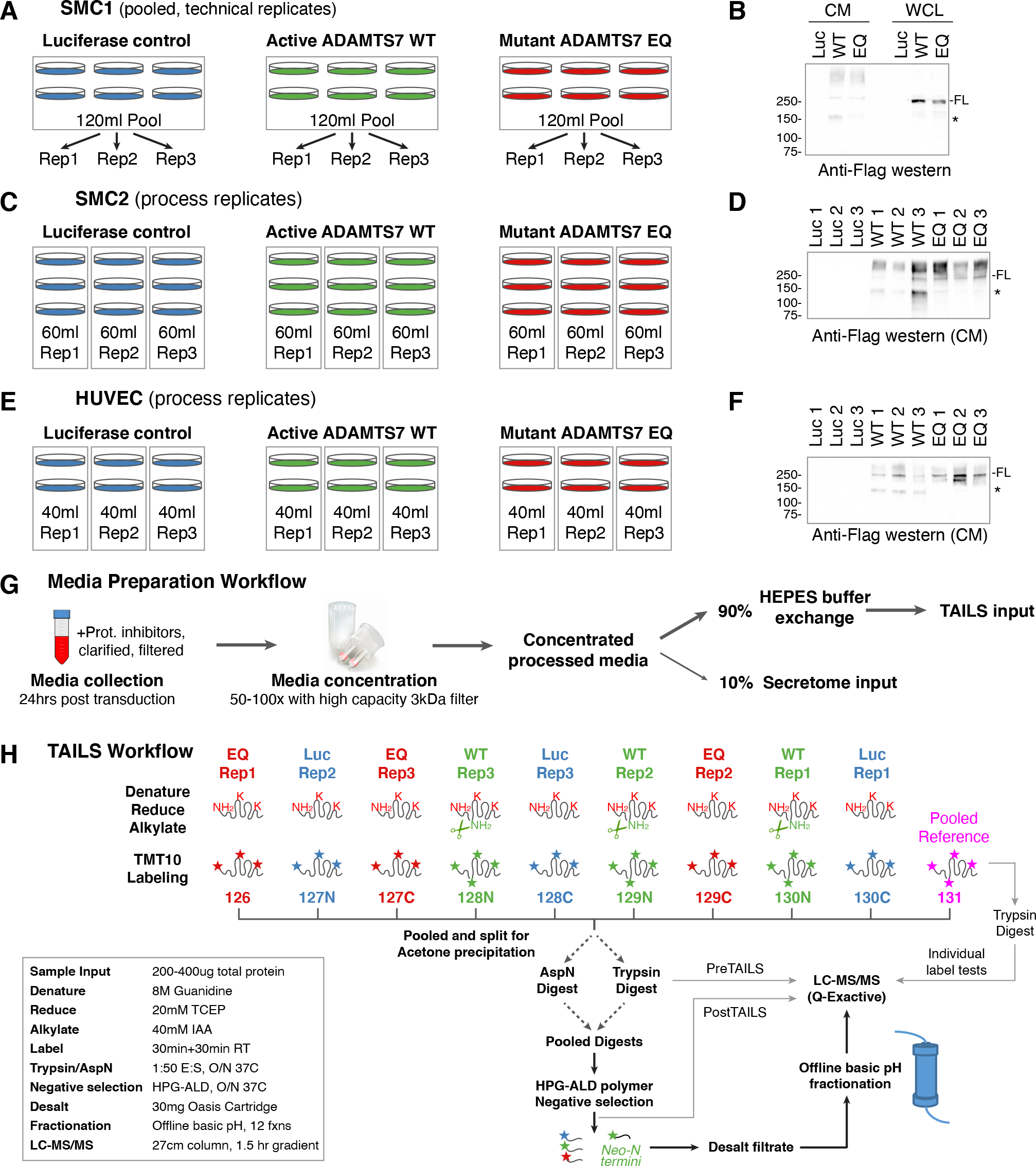
Proteomics sample input, media processing and TAILS workflow. Media collected from human coronary artery smooth muscle cells (SMC) or human umbilical vein endothelial cells (HUVEC) expressing control Luciferase (Luc), active mouse ADAMTS7 (WT) or catalytic mutant mouse ADAMTS7 E373Q (EQ) from three separate experiments. 20ml of media collected from each 15cm tissue culture dish. *A*, SMC1 media was pooled for each condition and split into technical replicates after concentration to generate technical replicates. *B*, expression of full-length (FL) ADAMTS7-3xFLAG constructs was verified in the conditioned media (CM) and whole cell lysate (WCL) by direct anti-Flag HRP western blot detection. * indicates mucin domain cleaved degradation product detected by c-terminal Flag tags. *C*, Replicates from SMC2 were collected from 3 dishes and processed separately. *D*, expression in the media from SMC2 replicates was verified by western blot. *E*, Replicates from HUVEC were collected from 2 dishes and processed separately. *F*, expression in the media from HUVEC replicates was verified by western blot. *G*, media preparation workflow for each replicate to generate input for total secretome and TAILS proteomics experiments. *H*, sample processing for TMT10 TAILS proteomics to identify neo-N-termini from the active ADAMTS7 enzyme condition. SMC1 TAILS experiment was digested with Trypsin only. SMC2 and HUVEC were digested with AspN or Trypsin before negative selection.

**Figure S2.**
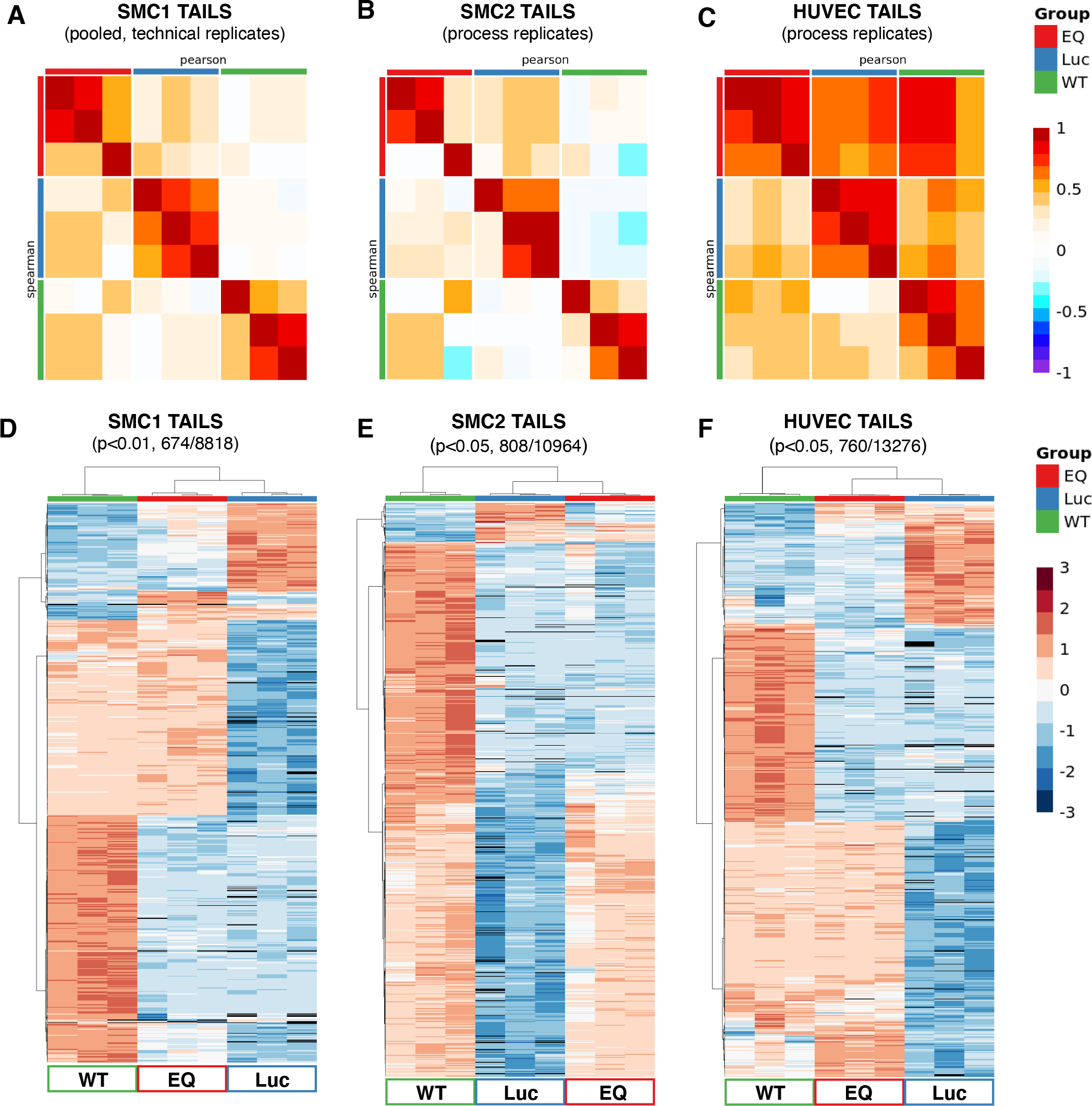
TAILS replicate correlation plots and heatmaps for regulated peptides. A-C, Similarity of TAILS experimental replicates analyzed by Pearson (linear relationships) and Spearman (monotonic relationships) correlation plots generated by Protigy. D-F, Heatmap cluster analysis of regulated peptides from TAILS experiments demonstrating greater clustering of EQ and Luc compared to WT replicates associated with ADAMTS7 activity.

**Figure S3.**
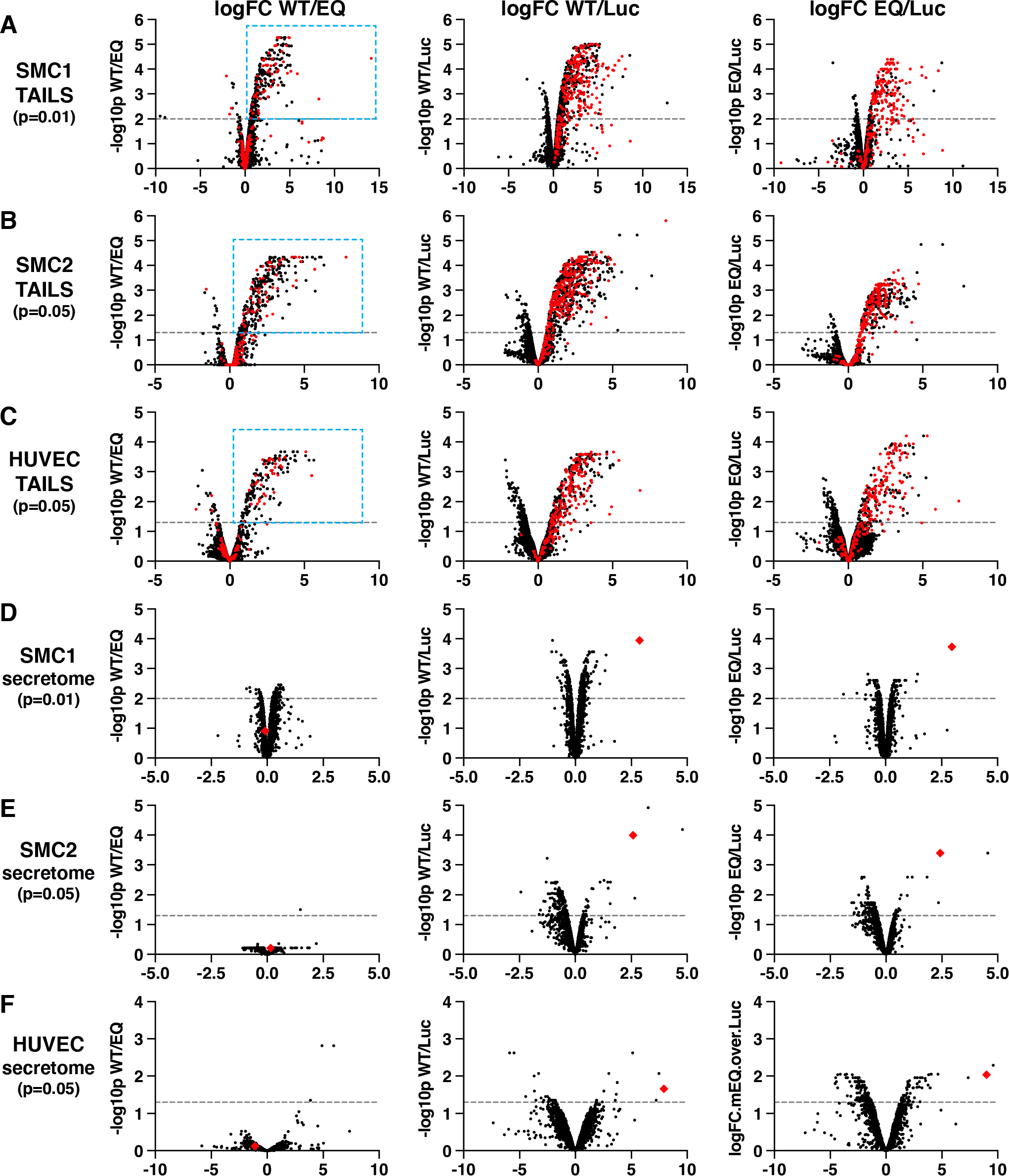
Volcano plots of regulated TAILS peptides and secretome regulated proteins. *A-C*, comparison of WT/EQ, WT/Luc and EQ/Luc regulated peptides from three independent TAILS experiments. *A*, TAILS SMC1 (p<0.01) *B*, TAILS SMC2 (p<0.05) *C*, TAILS HUVEC (p<0.05). Outlier logFC values with incomplete replicate data were excluded from some SMC1 volcano plots to keep similar data ranges. Mouse ADAMTS7 peptides are shown in red. Significant WT/EQ peptides with positive logFC are boxed in cyan and represent candidate substrate cleavage sites and ADAMTS7 auto-cleavage sites. *D-F*, comparison of WT/EQ, WT/Luc and EQ/Luc regulated proteins from the total secretome analysis. *D*, TAILS SMC1 (p<0.01) *E*, TAILS SMC2 (p<0.05) *F*, TAILS HUVEC (p<0.05). Mouse ADAMTS7 total protein (red diamonds) was significantly upregulated in the WT/Luc and EQ/Luc comparisons.

**Figure S4.**
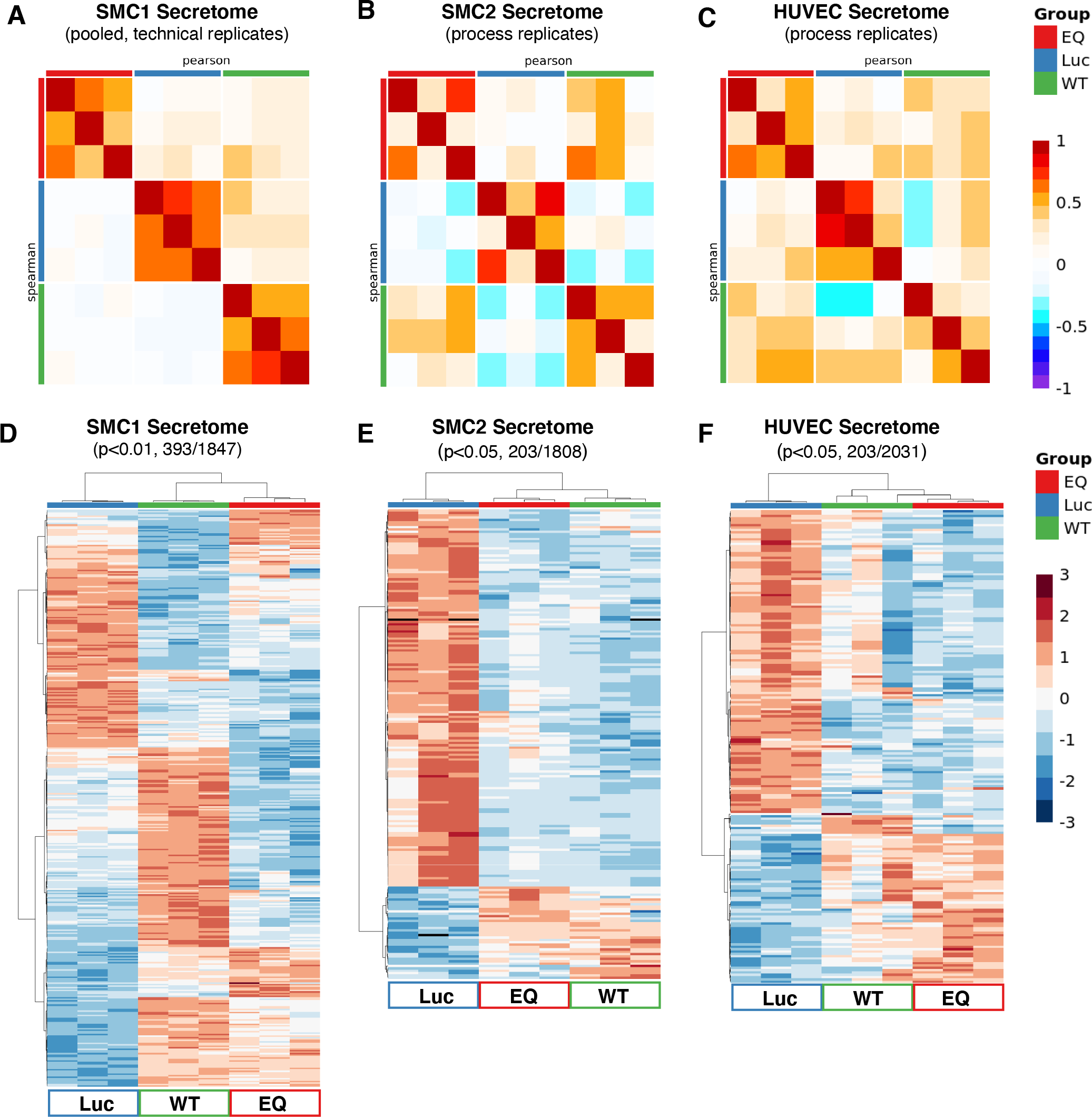
Secretome replicate correlation plots and heatmaps for regulated proteins. A-C, Similarity of secretome experimental replicates analyzed by Pearson (linear relationships) and Spearman (monotonic relationships) correlation plots generated by Protigy. For the total secretome analysis, a standard median normalization was used for the SMC1 technical replicates. SMC2 and HUVEC samples displayed greater diversity from biological replicates, therefore we applied median and median absolute deviation (M-MAD) to normalize the more variable secretome samples. D-F, Heatmap cluster analysis of regulated proteins from secretome experiments demonstrating greater clustering of WT and EQ compared to Luc replicates. The secretome heatmap dendrogram differs from the TAILS heatmap dendrogram and may be a product of Ad-ADAMTS7 expression rather than ADAMTS7 activity.

**Figure S5.**
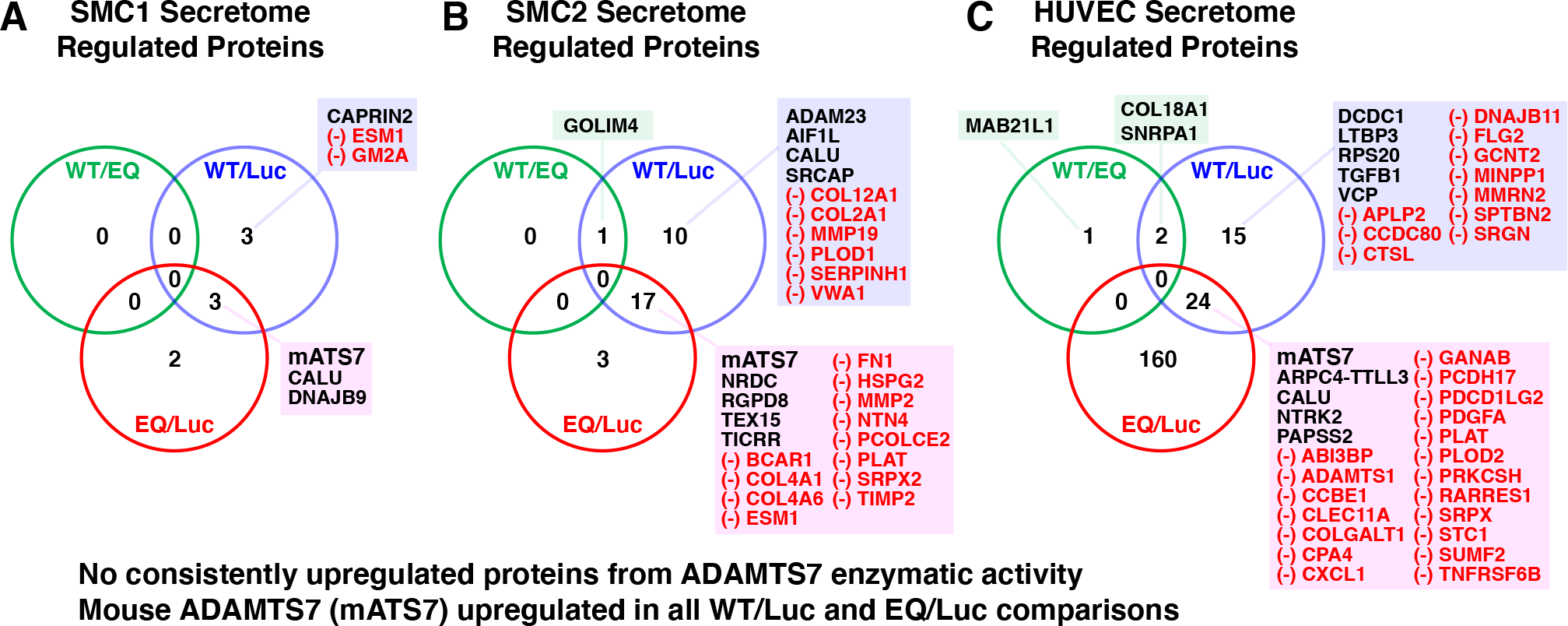
Analysis of overlapping regulated proteins from each secretome experiment. Comparison of regulated proteins within each ADAMTS7 secretome experiment. Significantly upregulated proteins (logFC >1) and down-regulated proteins (logFC < -1) shown in red. List of proteins regulated in EQ/Luc alone are not shown. A. SMC1 Venn diagram, B. SMC2 Venn diagram, C. HUVEC Venn diagram.

**Figure S6.**
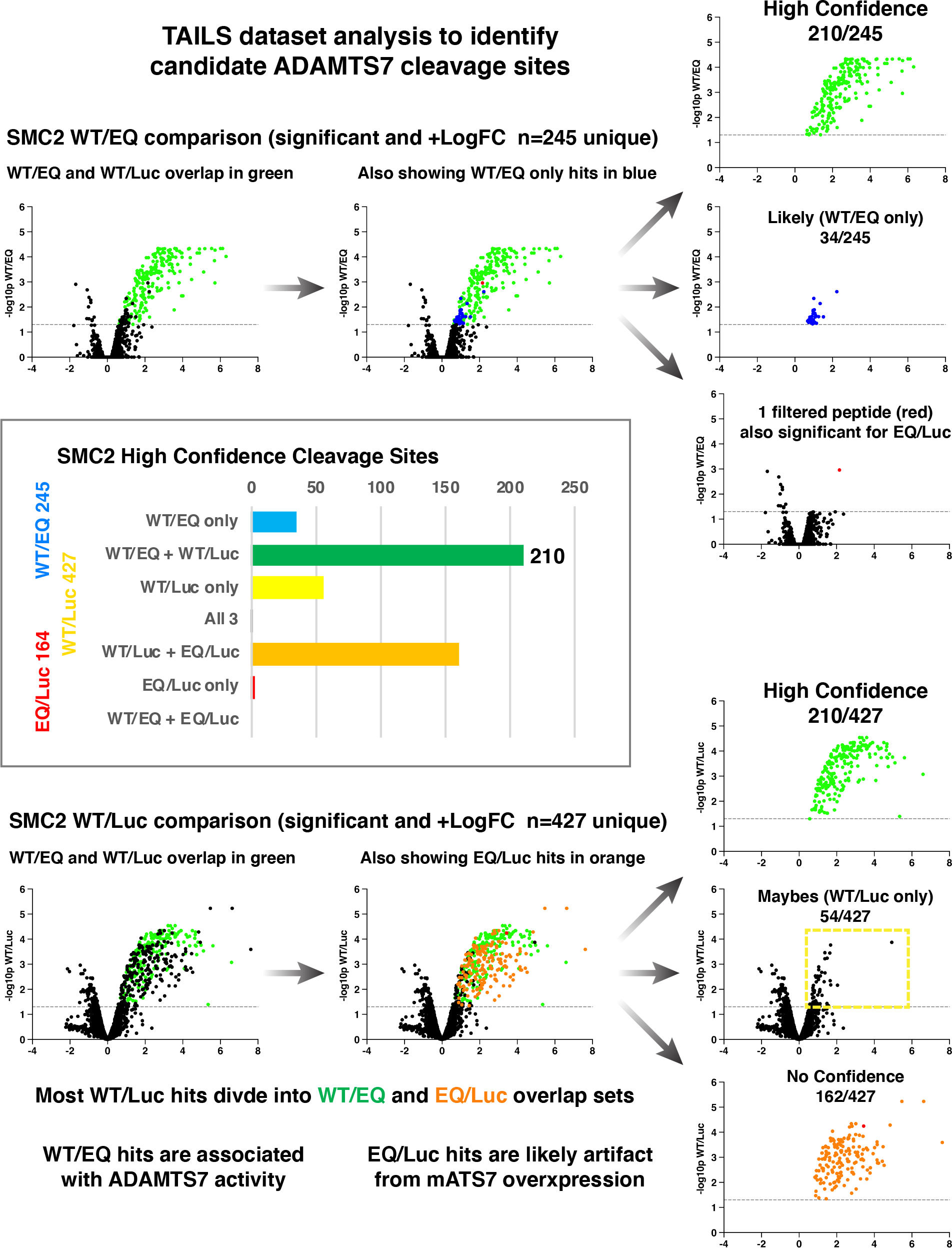
Candidate assessment from SMC2 WT/EQ and SMC2 WT/Luc TAILS comparisons after removal of auto-cleavage sites. Detailed breakdown of candidates from volcano plots shown in Figure 4B to illustrate the 210 high confidence substrate cleavage sites and remaining unqualified regulated peptides. Panel from Figure 5C is included to show the distribution of SMC2 regulated peptides. A majority of WT/EQ regulated peptides met all the criteria for high confidence substrate cleavage sites, while roughly half of the WT/Luc regulated peptides were categorized as high confidence. In the case of the SMC2 TAILS dataset, the WT/EQ only average logFC was 1.0 versus the WT/EQ high confidence average of logFC of 2.5. While a predominance of high confidence hits were present in the WT/EQ regulated peptides, the WT/Luc regulated peptides contained a mixture of overlap with the activity associated WT/EQ comparison and the EQ/Luc regulated peptides likely associated with artifact from ADAMTS7 overexpression. For simplicity numbers reflect unique substrate cleavage sites, however the volcano plots contain all regulated peptides including multiple identifications for ADAM9_69 and COL1A2_113 within the SMC2 TAILS dataset.

**Figure S7.**
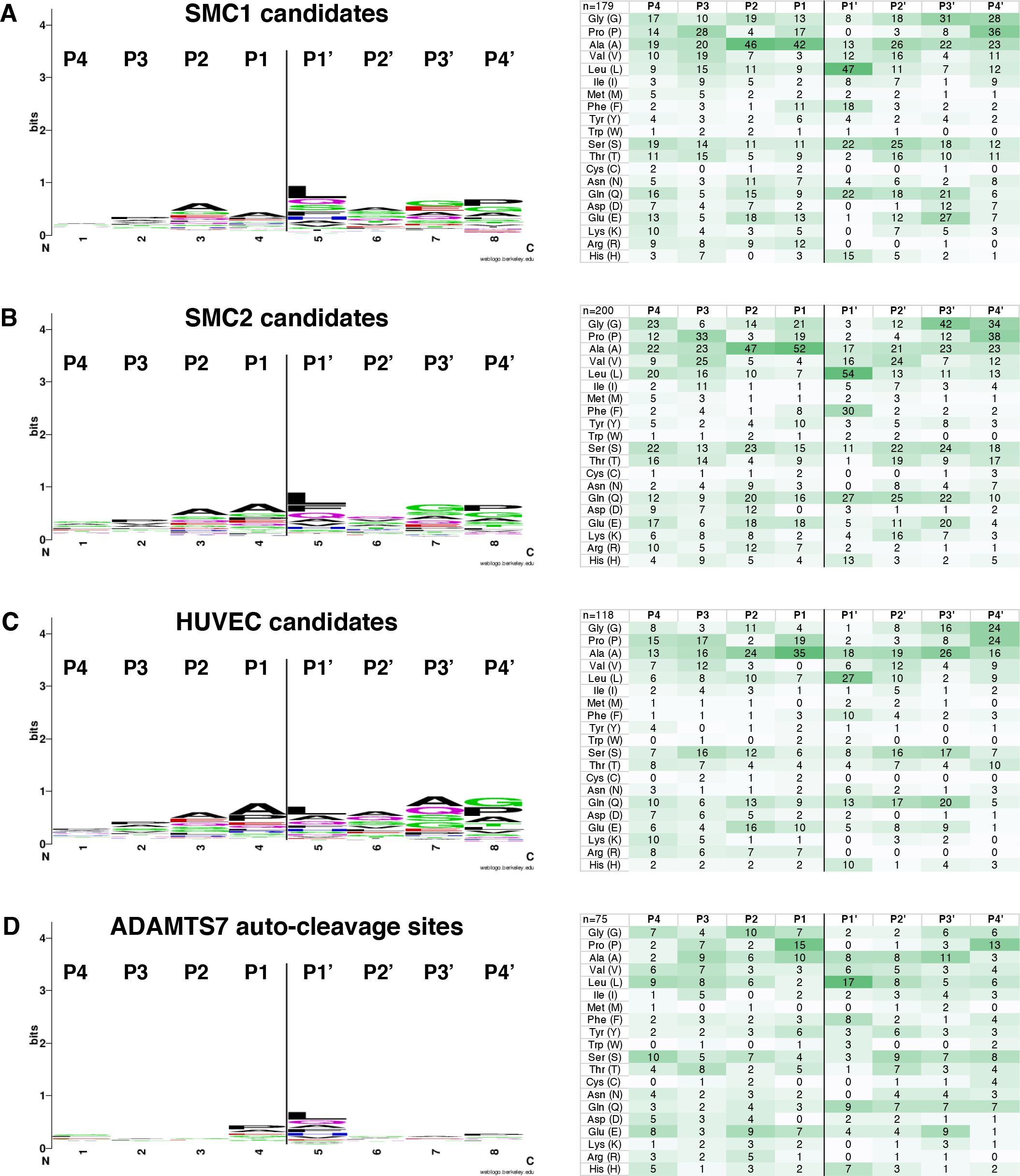
ADAMTS7 cleavage site specificity from TAILS experiments. A-D, Stringent cleavage site logo plots generated by WebLogo and amino acid counts for the TAILS high confidence candidate substrate cleavage sites and ADAMTS7 auto-cleavage sites. A, SMC1 candidates (p<0.01, +FC>1, n=179), B, SMC2 candidates (p<0.05, +FC>1, n=200), C, HUVEC candidates (p<0.05, +FC>1, n=118) and D, all unique ADAMTS7 auto-cleavage sites (n=75).

**Figure S8.**
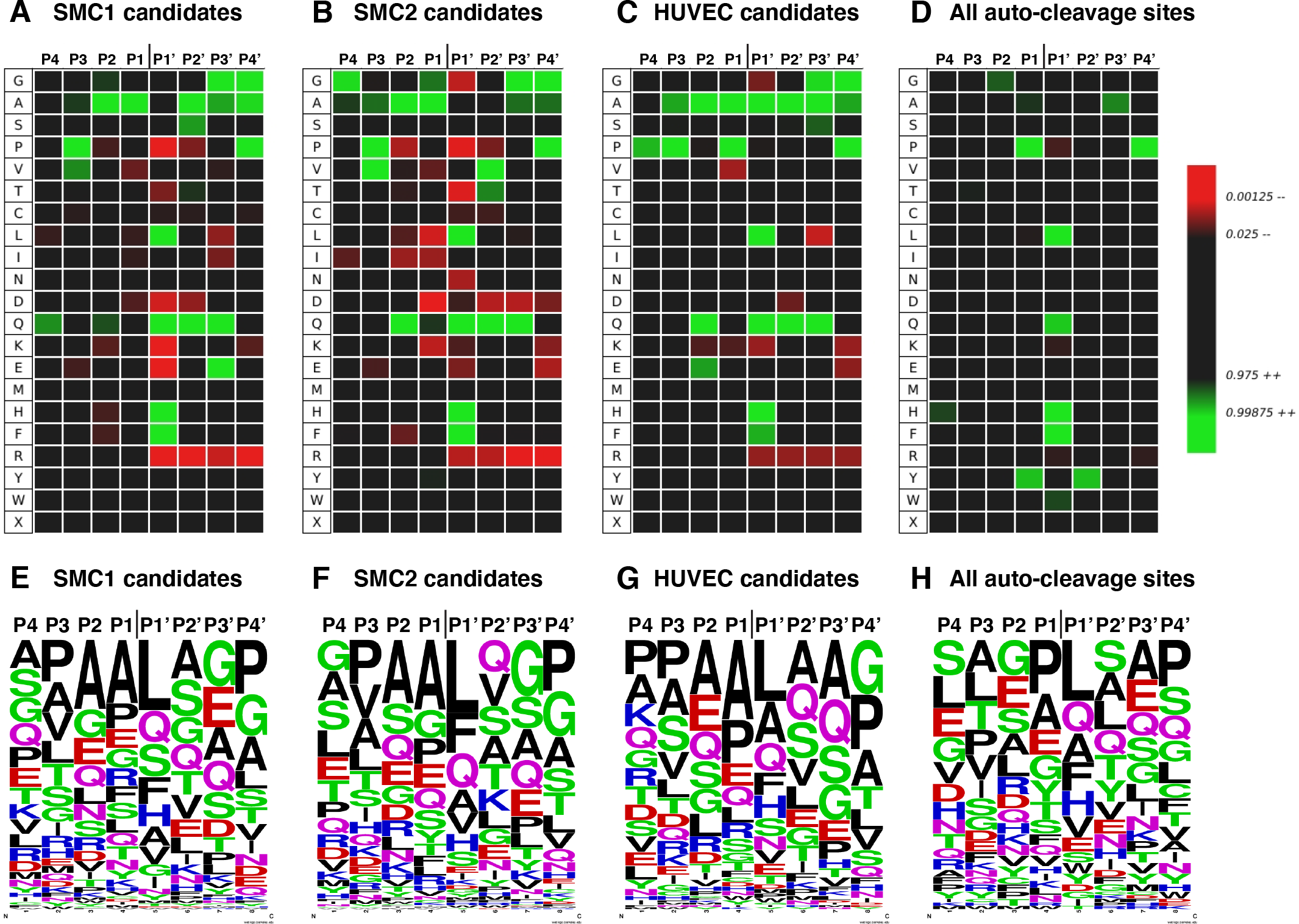
TAILS high confidence substrate site specificity compared with ADAMTS7 auto-cleavage sites. *A-D*, TAILS candidate substrate cleavage sites and ADAMTS7 auto-cleavage sites analyzed using iceLogo to generate heatmaps adjusted for the abundance of each amino acid in humans. *A*, SMC1 candidate heatmap (p<0.01, +FC>1, n=179), *B*, SMC2 candidate heatmap (p<0.05, +FC>1, n=200), *C*, HUVEC candidate heatmap (p<0.05, +FC>1, n=118). *D*, the heat map including all unique ADAMTS7 auto-cleavage sites (n=75) resembles the TAILS high confidence substrates. *G-H*, Amino acid frequency plots generated by WebLogo showing the similar distribution between experiments, however no amino acid was present more than 30% at any given position at the cleavage site.

**Figure S9.**
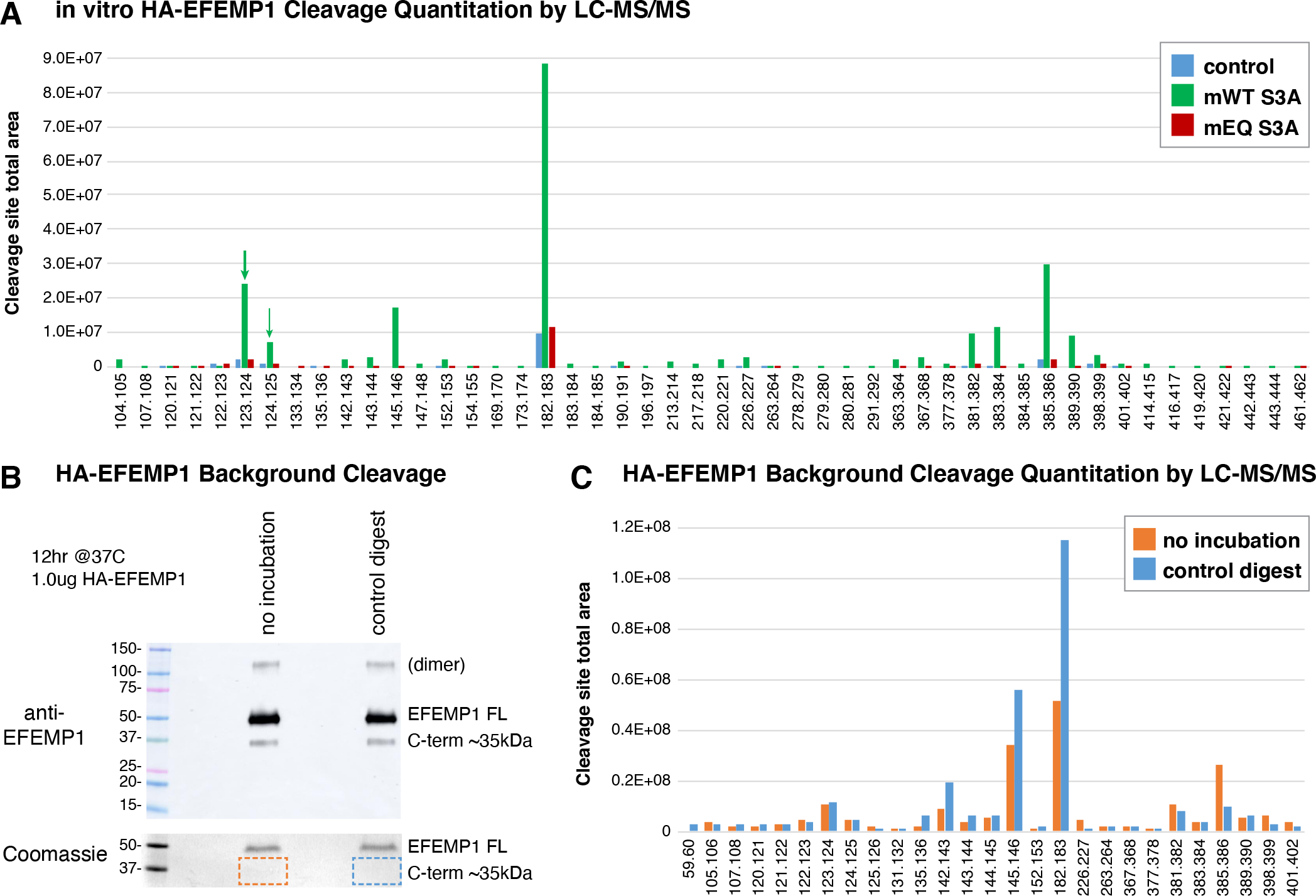
Purified HA-EFEMP1 in vitro cleavage and background cleavage. A, quantitation of semi-tryptic or semi-chymotryptic HA-EFEMP1 peptides from the ADAMTS7 in vitro cleavage assay. A subset of this data at the atypical EGF1 repeat was presented in Figure 7. ADAMTS7 specific cleavage sites at 123.124 and 124.125 are indicated by green arrows. B, background cleavage in purified HA-EFEMP1 within the atypical EGF1 repeat in the absence of enzyme shown by western blot under non-reducing conditions. A region below the full-length HA-EFEMP1 was excised from a parallel Coomassie stained non-reducing gel and sent off for LC-MS/MS identification of EFEMP1 peptides. C, quantitation of semi-tryptic or semi-chymotryptic peptides showing background cleavage from the commercial purified HA-EFEMP1 protein. Background cleavage at positions 123.124 and 124.125 were detected and did not increase in abundance after overnight digestion in control conditions.

