## supplemental Table S1 for "TAILS identifies candidate substrates and biomarkers of ADAMTS7, a therapeutic protease target in coronary artery disease"

**Supplementary Table 1 Quantitation statistics of the large scale SMC and HUVEC TAILS experiments.** TMT labeled peptides filtered for redundancy, species, and DeltaForwtoRev > 0. Peptides with unmodified or acetylated N-termini containing internal TMT labeled lysine residues were also included in the analysis.

| | **SMC1** | | **SMC2** | | **HUVEC** | |
|---|---|---|---|---|---|---|
| | Peptides | Proteins | Peptides | Proteins | Peptides | Proteins |
| Quantified TMT labeled peptides | **8818** | **3152** | **10964** | **3579** | **13276** | **3826** |
| Peptides N-terminally labeled with TMT | 7325 | 2618 | 6482 | 2349 | 7716 | 2685 |
| Peptides with unmodified N-termini | 926 | 483 | 3579 | 1231 | 4873 | 1567 |
| N-terminally acetylated peptides | 567 | 529 | 903 | 831 | 687 | 636 |
