## supplemental Table S4 for "TAILS identifies candidate substrates and biomarkers of ADAMTS7, a therapeutic protease target in coronary artery disease"

**Supplementary Table 4 Summary of regulated peptides in SMC and HUVEC TAILS experiments.** Significantly regulated quantitated peptides from WT/EQ, WT/Luc and EQ/Luc comparisons presented from each TAILS experiment. Positive Log fold change (FC+) and negative Log fold change (FC-) regulated peptides indicate significantly upregulated or downregulated peptides from each comparison. After removal of ADAMTS7 peptides, the candidate FC+ represent potential ADAMTS7 substrate cleavage sites. High confidence candidates were present in both WT/EQ and WT/Luc comparisons from each TAILS experiment.

| | **8818** | **SMC1 p<0.01** | | **10964** | **SMC2 p<0.05** | | **13276** | **HUVEC p<0.05** | |
|---|---|---|---|---|---|---|---|---|---|
| | **WT/EQ** | **WT/Luc** | **EQ/Luc** | **WT/EQ** | **WT/Luc** | **EQ/Luc** | **WT/EQ** | **WT/Luc** | **EQ/Luc** |
| **Significant** | **312** | 646 | 217 | **314** | 755 | 333 | **213** | 524 | 349 |
| All LogFC+ | 297 | 590 | 194 | 302 | 681 | 321 | 191 | 376 | 251 |
| All LogFC- | 15 | 56 | 23 | 12 | 74 | 12 | 22 | 148 | 98 |
| ADAMTS7 FC+ | **57** | 201 | 133 | **54** | 250 | 157 | **36** | 157 | 136 |
| ADAMTS7 FC- | 3 | 0 | 0 | 1 | 0 | 0 | 3 | 1 | 0 |
| Candidates FC+ | **240** | 389 | 61 | **248** | 431 | 164 | **155** | 219 | 115 |
| Candidates FC- | 12 | 56 | 23 | 11 | 74 | 12 | 19 | 147 | 98 |
| Dup Cand FC+ | 2 | 3 | 0 | 3 | 4 | 0 | 1 | 1 | 1 |
| Novel Cand FC+ | **238** | 386 | 61 | **245** | 427 | 164 | **154** | 218 | 114 |
| High confidence | **207** | | | **210** | | | **127** | | |
